# Single-nucleus multiomics unveils malignant cellular states, regulatory architectures and microenvironmental reorganization across the G-CIMP epigenomic transition in IDH-mutant glioma

**DOI:** 10.64898/2026.06.05.730437

**Authors:** Grayson A. Herrgott, Luciano Garofano, Natalia Silva Morosini, Bogdan Done, Christopher Powell, Ana deCarvalho, Laura Hasselbach, Andrea Transou, Ian Lee, Tobias Walbert, James Snyder, Anna Lasorella, Ana Valeria Castro, Antonio Iavarone, Houtan Noushmehr

## Abstract

IDH-mutant gliomas stratified by glioma CpG island methylator phenotype (G-CIMP) into High (GCH) and Low (GCL) exhibit markedly divergent clinical outcomes, yet cellular and regulatory determinants of this distinction remain incompletely defined. Integrating single-nucleus RNA-and ATAC-sequencing across 18 tumor specimens from 10 patients, we resolved six malignant cellular states whose differential enrichment across G-CIMP strata delineates the GCL epigenomic transition. GCL tumors were enriched for independently prognostic Mesenchymal and Mitotic Proliferative states, driven by convergent E2F, MYC, MEF2, and NFI-family regulatory networks confirmed across chromatin, histone, and transcriptomic modalities. Pseudotime trajectory inference revealed a multifurcating developmental model from an Astrocytic-like origin, with GCL tumors gaining preferential access to proliferative and mesenchymal endpoints. A reorganized immune microenvironment and candidate therapeutic axes including *CDK4/6*-*E2F*, *MYC*/*BET*, *KIF11*, and NOTCH, potentially combinable with vorasidenib as an epigenomic backbone provide a translationally actionable framework for intercepting GCL progression in IDH-mutant glioma.

## Introduction

IDH-mutant diffuse gliomas produce the oncometabolite D-2-hydroxyglutarate (2-HG), which competitively inhibits α-KG-dependent dioxygenases including TET methylcytosine hydroxylases, inducing widespread DNA hypermethylation and the glioma CpG island methylator phenotype (G-CIMP) ^1,2^. G-CIMP was first described as a robust, reproducible marker of IDH-mutant biology and a powerful complement to histopathological assessment ^3^. Integrative analyses have shown IDH-mutant tumors further stratify by histomolecular lineage (e.g., oligodendroglioma with 1p/19q codeletion, astrocytoma with ATRX/TP53 alteration); G-CIMP meanwhile captures shared epigenomic consequences across lineages ^4–6^. A central advance arising from methylation-based classification is recognition of a clinically consequential epigenomic axis; G-CIMP-high (GCH) tumors reside in a globally hypermethylated landscape corresponding to differentiated cellular programs, relative genomic stability, and favorable outcomes ^4,7^. In contrast, G-CIMP-low (GCL) tumors exhibit partial loss of DNA methylation, widespread enhancer remodeling, and derepression of developmental and stem-like transcriptional networks ^7,8^. Clinically, GCL status is strongly associated with uncharacteristically higher histological grade, inferior overall survival, and recurrent disease, yet its mechanistic and cellular underpinnings remain incompletely characterized ^9,10^.

Prior bulk transcriptomic and epigenomic profiling has highlighted genome-wide contrasts between GCH and GCL, however, cannot resolve which specific malignant cell states, chromatin accessibility programs, or microenvironmental interactions drive GCL-associated aggressiveness. Single-cell studies in IDH-mutant gliomas have revealed developmental hierarchies spanning astrocyte-like and oligodendrocyte precursor-like populations, stem-like progenitors, and mesenchymal programs, alongside dynamic immune and stromal compartments ^11–13^. And yet, no study has systematically interrogated these features across the G-CIMP epigenomic axis. Potential interventions, such as brain-penetrant IDH inhibitors (e.g., vorasidenib) may delay GCL transition by reducing 2-HG, making cellular resolution of the GCL state translationally urgent ^2,14^. Yet, the extent and durability of epigenomic reprogramming required for reversal or prevention of GCL-associated regulatory states, is not fully understood. We endeavoured to define how G-CIMP evolution shapes malignant cellular diversity, chromatin accessibility programs, and tumor-microenvironment interactions, providing mechanistic insight into the GCL phenotype and identifying convergent vulnerabilities for therapeutic interception.

## Results

### Cohort description & experiment detailing

We generated a single-nucleus RNA- and ATAC-sequencing (snRNA/ATAC-seq) multiomic dataset from 18 tumor samples collected from 10 patients, including samples from initial (n=9) and recurrent (first recurrence: n=4; second recurrence: n=5) time points (**Supplementary File S1**). Samples were further stratified by glioma CpG island methylator phenotype (G-CIMP) status, defined by widespread CpG hypermethylation based on prior bulk DNA methylation profilings.

Following quality control, 20,967 cells were retained (**Supplementary Figure S1a**). Malignant (n=10,366) or non-tumorous (n=4,602) cells were distinguished through inferred copy number alterations (CNAs) and graph-based clustering trends; 5,999 cells with ambiguous CNV profiles were excluded (**Figure 1a-c**; **Supplementary Figure S1b**).

**Figure 1.**
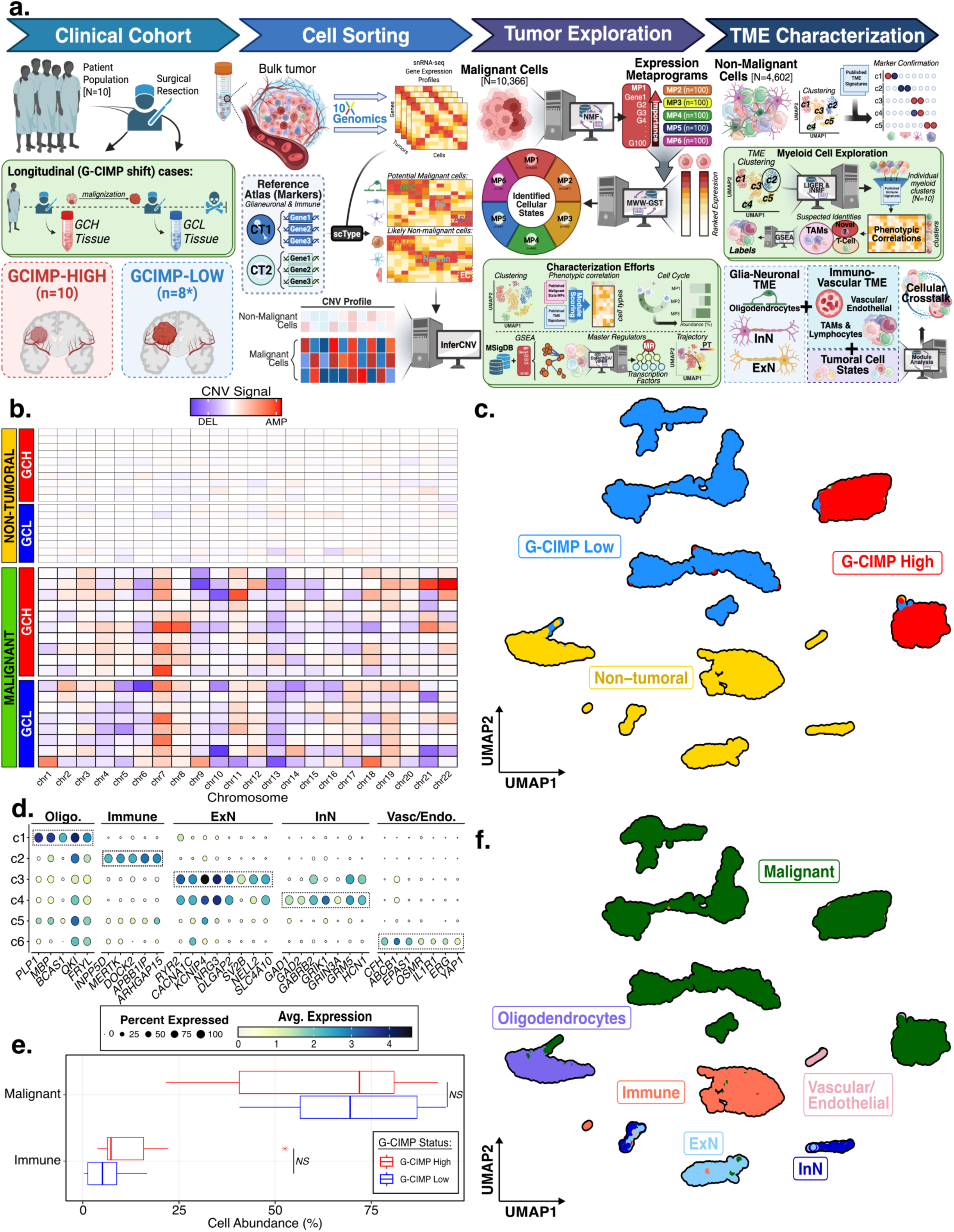
Study overview and cohort detailing. **a)** Experimental workflow schematic. **b)** Sample-level copy number signals, with samples (rows) segregated by G-CIMP. Values reflect the average signal across all cells classified as malignant, or nontumor. **c)** Uniform manifold approximation and projection (UMAP) for dimension reduction based on gene expression of the 14,968 cells in our cohort. **d)** Expression-based dot plot of canonical cell type markers across non-tumor clusters. Dots are colored by the average expression value, and sized by the percentage of cells expressing each marker. Notes: Oligo: oligodendrocyte; ExN: Excitatory Neurons; InN: Inhibitory Neurons; Vasc/Endo: vascular/endothelial. **e)** Boxplot displaying cell abundance across samples, segregated by G-CIMP. Note: NS: Non-significant. **f)** UMAP for dimension reduction using gene expression of the 14,968 cells in our cohort.

Harmony-corrected UMAP visualization revealed transcriptional heterogeneity across the malignant population, with separation by G-CIMP methylation status and intra-GCL heterogeneity, consistent with known transcriptional diversity of IDH-mutant gliomas (**Figure 1c**) ^15^. Non-tumoral cells were classified into five robust cell clusters: oligodendrocytic, excitatory and inhibitory neuronal (ExN, InN), vascular/endothelial, and immune (**Figure 1d-f**; **Supplemental Figure S1c-e**) ^16,17^. No significant differences in non-tumoral cell abundance were observed across G-CIMP categorizations or WHO grade.

### Definition of malignant cellular states across IDH-mutant glioma

Malignant cellular states were defined using Non-negative Matrix Factorization (NMF), yielding six distinct, robust, and non-redundant transcriptional metaprograms (MP1-6) shared across samples (**Figure 2a, Supplementary File S2**) ^18^. Malignant cells were assigned to cellular states according to their highest MP module score (*AddModuleScore*); low-confidence cells were left unassigned (**Figure 2b, Supplementary Figure S2a**) ^19^. State identities were validated through correlation analyses against normal-brain cell populations and previously established malignant cell states from IDH-wildtype and IDH-mutant gliomas (**Figures 2c-2d**, **Supplementary Figure S2b**) ^20–27^. Cell cycle phases were assigned using ccAFv2 framework (**Figures 2e-f**, **Supplementary Figure S2d-e**) ^28^. To identify the transcriptional regulatory logic underlying each cellular state, we applied the VIPER algorithm to infer transcription factor (TF) activity from coordinated target gene expression (**Figure 2g**; **Supplementary Figure S2g**) ^29^.

**Figure 2.**
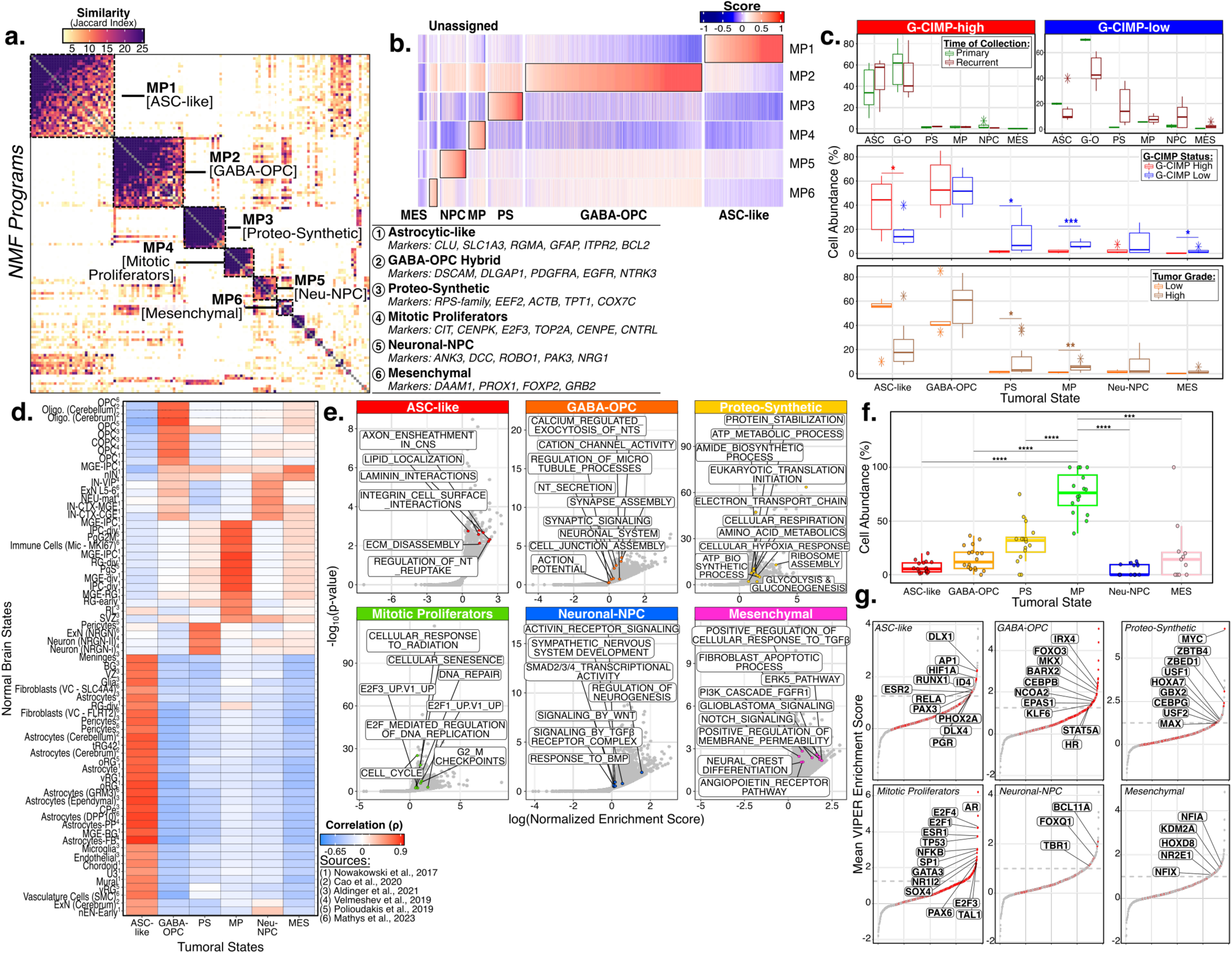
Malignant state definition in IDHmut glioma. **a)** Jaccard similarity indices of robust and non-redundant NMF programs (n =131), ordered according to clustering and grouped into metaprograms (MPs). Key markers for each of the named metaprograms are listed. **b)** Heatmap depicting module scores of metaprograms across cells. **c)** Boxplot displaying cell abundance across samples for cellular states, split by (i) combinatory G-CIMP and Time of Collection (ii) G-CIMP, and (iii) WHO Grade. **d)** Correlative heatmap relating expression of published cell type signatures and malignant metaprograms. **e)** Scatter plots depicting the enrichment of selected pathways. Y-axis: log-scaled significance (p-value); x-axis: normalized enrichment scores. **f)** Boxplot displaying actively cycling cell abundance per sample for cellular states. **g)** Rank order plots for transcription factors (TFs) in each cell state. Red dots indicate statistically significant TFs which are representative and uniquely enriched. Notes: ASC-like: Astrocytic-like; PS: Proteo-Synthetic; MP: Mitotic Proliferative; Neu-NPC: Neuronal-NPC; MES: Mesenchymal. Boxplots: *: p<0.05; **: p<0.01, ***: p<0.001; ****: p<0.0001.

MP1 (Astrocytic-like), displayed expression of canonical astrocytic markers (*GFAP*, *CLU, APOE, SLC1A2/3, LIFR, BMPR1A/B*), and markers indicative of CNS microenvironment action (*DCLK1/2, ROBO2, SEMA6A, ARHGEF4*)(**Figure 2a**). While specific for GCH, correlations demonstrated strong relation with normal-brain astrocytic signature sets and previously-defined malignant astrocytic-like programs (**Figure 2c-d**, **Supplementary Figure S2b**). Ontological enrichments mirrored astrocytic-like and reactive astrocyte programs (axon ensheathment, lipid localization)(**Figure 2e**, **Supplementary Figure S2e**). MP1 cells were predominantly G0/G1 (**Figure 2f**, **Supplementary Figure S2d**). Inferred master regulators included developmental homeobox factors and *HIF1A*, consistent with metabolic stress adaptation in the hypoxic tumor microenvironment (**Figure 2g, Supplementary Figure S2g**) ^30–33^.

MP2, a broadly distributed (40-70%) GABA-ergic neuronal-oligodendrocyte precursor cell (OPC) malignant hybrid state (hereafter, GABA-OPC), exhibited expression of canonical OPC (*PDGFRA, PTPRZ1, SOX6, VCAN*), core neuronal and GABA-ergic markers (*GAD1, MAP2, NXPH1, GRIA2*)(**Figure 2a**). Correlative analysis demonstrated strong association with nontumoral OPC and inhibitory neuronal signatures, in addition to reported malignant OPC programs (**Figure 2c-d**, **Supplementary Figure S2b**). Enriched ontologies encompassed neuronal processes, GABAergic transmission, and extracellular matrix functions (**Figure 2e**, **Supplementary Figure S2e**). MP2 cells were similarly predominantly G0/G1 (**Figure 2f**, **Supplementary Figure S2d**). MRs *FOXO3* and CEBPB collectively supported an uncommitted, plastic progenitor identity (**Figure 2g, Supplementary Figure S2g**) ^34,35^.

MP3 (Proteo-synthetic) was characterized by high expression of ribosomal biogenesis and core translation machinery (*RPL*/*RPS* subunits), translation elongation factors (*EEF1A1*, *EEF2*, *NACA*), and oxidative phosphorylation components (*ATP5F1E*, *COX7C*)(**Figure 2a**) ^36–38^. While specific for higher-grade and GCL tumors, we observed correlations with the malignant ribosomal program identified in Nomura et al. (**Figure 2c-d**, **Supplementary Figure S2b**) ^39^. Enriched ontologies spanned energy metabolism (electron transport chain, glycolysis) and translation (protein stabilization, amino acid metabolism)(**Figure 2e**, **Supplementary Figure S2e**). MP3 cells were preferentially situated in active G2/M and S-phases, and master regulator *MYC*, alongside translational modulators *MAX* and *USF1/2*, were consistent with sustained protein production demand (**Figure 2f-g**, **Supplementary Figure S2d&g**) ^40–42^.

MP4 (Mitotic Proliferative) expressed canonical mitotic markers (*CIT*, *CENPK*, *TOP2A*, *CENPE*), as well as cell cycle regulators (*E2F3)*(**Figure 2a**). This state was GCL and high-grade enriched, and displayed strong correlation with intermediate progenitor cells (IPC) and previously defined proliferative progenitor programs (**Figure 2c-d, Supplementary Figure S2b**). A confluence of cellular survival, cell cycle and DNA maintenance ontologies produced a robust rendering of this highly proliferative identity (**Figure 2e, Supplementary Figure S2e**). We observed predictable G2/M and S-phase concentrations, and identified master regulators including E2F-family transcription factors (*E2F1/3/4*) and supporting DNA replication checkpoint regulators (**Figure 2g, Supplementary Figure S2g**) ^43,44^.

MP5, a neuronal-committed neural progenitor cell (NPC)-like state (hereafter, Neuronal-NPC), expressed core neuronal identity and neurodevelopmental hallmarks (*SOX11*, *MYT1L*), alongside committed progenitor markers (*DAAM1*, *PROX1*)(**Figure 2a**). Appreciable relation to nontumoral mature and ganglionic eminence neuronal programs, as well as malignant NPC entities, was observed (**Figure 2d; Supplementary Figure S2d**). Ontologies encompassed neuronal differentiation, dendritogenesis, axonogenesis, and synaptic plasticity(**Figure 2e, Supplementary Figure S2e**). Cells were predominantly positioned within the ‘inactive’ G0/G1 phase of the cell cycle (**Figure 2f**, **Supplementary Figure S2d**). Master regulators *BCL11A* and *TBR1*, consistent with post mitotic neuronal commitment factors and active cortical neurogenesis programs, were identified (**Figure 2g, Supplementary Figure S2g**) ^45,46^.

MP6 (Mesenchymal) expressed core mesenchymal transcriptional regulators and fate control markers (*MEF2A*, *MEF2C*, *FLI1*, *ETV6*), and hallmarks of mesenchymal motility (*ARHGAP12*/*15*, *DOCK2*/*4*/*8*, *SRGAP2*)(**Figure 2a**). This state was uniquely enriched in GCL tumors and showed no significant correlation with any previously defined malignant cellular state (**Figure 2c, Supplementary Figure S2b**). Ontological networks revealed consolidation into TGFβ-related, growth-factor driven motility/survival and state maintenance/plasticity structures (**Figure 2e, Supplementary Figure S2e**). In congruence with a mesenchymal identity, these cells occupied a primarily G0/G1 ‘inactive’ state (**Figure 2f**, **Supplementary Figure S2d**). NFI-family master regulators (*NFIA*, *NFIX*), alongside *HOXD8*, supported a developmentally permissive but non-neural regulatory program, influencing mesenchymal behaviors such as cellular migration and invasion (**Figure 2g, Supplementary Figure S2g**) ^47–49^.

### Refinement of neuronal-related malignant cellular states through multi-tiered comparison efforts

To further distinguish GABA-OPC (MP2) and Neuronal-NPC (MP5) populations, which demonstrated overlapping correlations to non-tumoral neuronal cell types, we applied serial characterization approaches. MP5 cells showed preferential expression of excitatory neurotransmitter receptors (nicotinergic acetylcholine, NMDA), whereas GABA-OPC cells exhibited diverse enrichment of both inhibitory and excitatory receptor families (e.g., AMPA, GABA-A/B, Kainate; **Figure 3a, Supplementary Figure S3a**). Principal component analysis (PCA) across inhibitory (InN) and excitatory (ExN) neuron, and OPC transcriptional module scores demonstrated orthogonal separation of the two states along PC1 and PC3 (>70% of total variance), with InN signatures correlating preferentially with GABA-OPC over Neuronal-NPC expression (**Figure 3b**; **Supplementary File S2**) ^50–55^. Additional correlative testing revealed InN signatures correlated well with the GABA-OPC metaprogram, statistically distinguished from Neuronal-NPC correlations (Wilcoxon rank-sum test, p=1.3e-06; **Figure 3c**).

**Figure 3.**
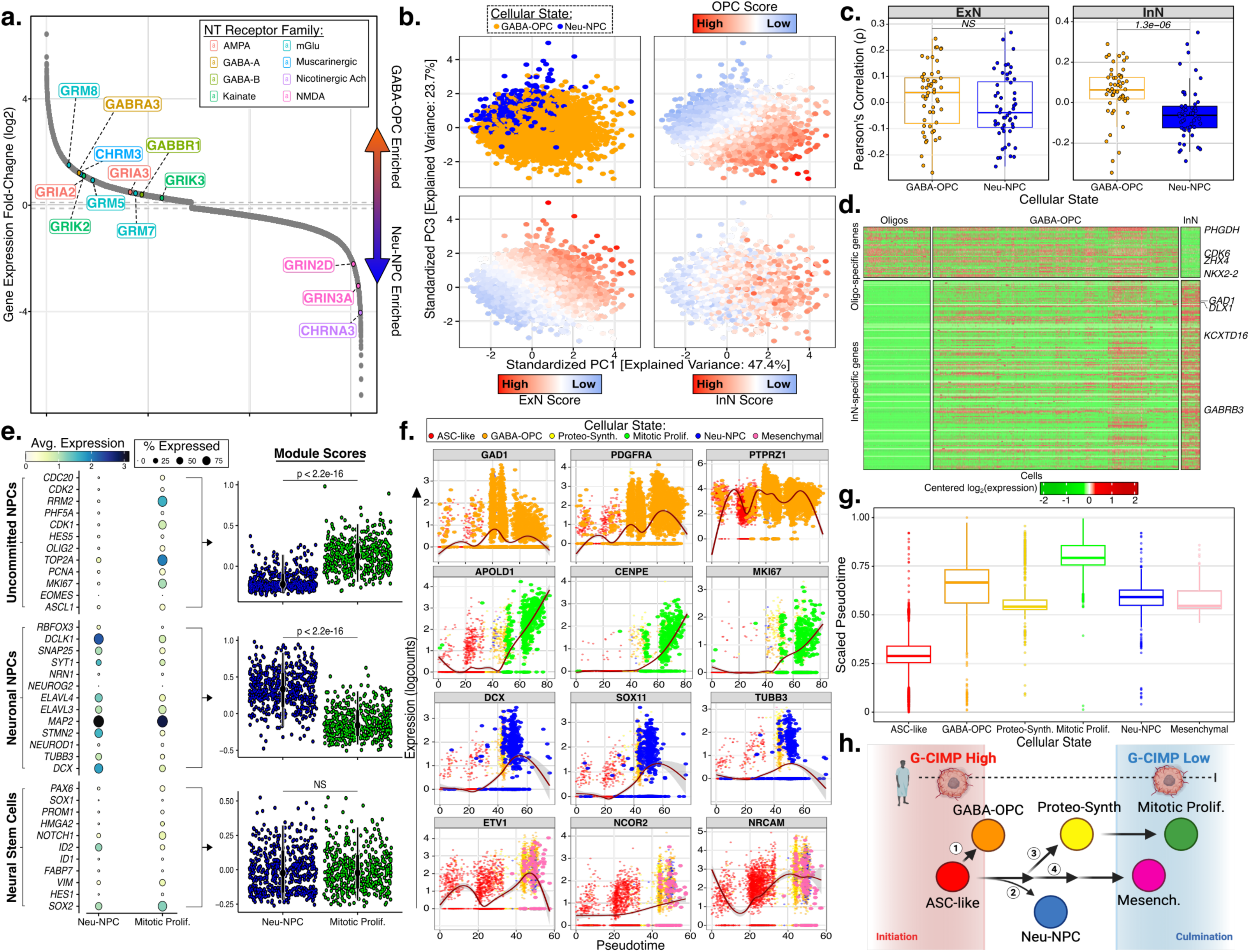
Malignant state discrimination and trajectory. **a)** Rank order plot displaying expression shifts between cells belonging to GABA-OPC and Neuronal-NPC populations. Notes: NT: neurotransmitter. **b)** Principal Component Analysis (PCA) using scores derived from inhibitory or excitatory neurons (InN, ExN), or oligodendrocyte precursor (OPC) signatures across GABA-OPC or Neu-NPC cells. Cells are colored according to their metaprogram assignment or signature score. **c)** Boxplot displaying Pearson correlation coefficients for metaprograms and ExN or InN signatures. **d)** Expression-based heatmap of the gene signatures which distinguish nontumoral cell types across both GABA-OPC and nontumoral cells. **e)** (Left) Dotplot displaying the expression of markers related to uncommitted/multipotent neural precursor cells (NPCs), neuronal-committed NPC, and neural stem cells across malignant cellular states. (Right) Geyser plots displaying related module scores. **f)** Scatterplots displaying temporally relevant marker expression. Cells are colored by cellular state. Projections of gene behavior are presented, with 95% confidence intervals. **g)** Boxplot displaying the dispersion of scaled pseudotime across malignant cellular states. **h)** Developmental trajectory schematic.

At the single-cell resolution, GABA-OPC cells simultaneously co-expressed canonical oligodendrocytic (*NKX2-2*, *CDK6*, *PHGDH*) and GABAergic neuronal (*GAD1*, *DLX1*, *GABRB3*) markers, with additional enrichment of OPC hallmarks (*PDGFRA*, *OLIG2*) and GABA-lineage markers (*GAD1*, *ABAT*, *SCN3A*)(**Figure 3d**, **Figure S3b-e**).

Neuronal-NPC cells showed specific enrichments for neuronal-committed NPC markers (*DCLK1*, *DCX*) and significantly higher neuronal-committed NPC module scores relative to the Mitotic Proliferative state (Wilcoxon rank-sum test, p<2.2×10^-16^; **Figure 3e**, **Supplementary File S2**) ^56–62^. These findings, alongside neuronal correlations and progenitor cell ontologies (Wnt signaling, neural precursor proliferation), confirm MP5 as a neuronal-committed progenitor state (**Supplementary Figure 3f**).

### Pseudotime trajectory defines a multifurcating developmental model from a GCH Astrocytic-like origin

To explore developmental relationships among the six states, we applied Slingshot (v2.16) following Harmony-based correction of inter-sample transcriptional variation ^63,64^. Expression dynamics along each developmental branch were characterized using generalized additive models through tradeSeq (v1.22.0) (**Figure 3f-h, Supplementary Figure S3g**) ^65^. Four developmental trajectories were identified from a shared Astrocytic-like origin, consistent with this state representing the most differentiated and developmentally proximal identity.

Along the first trajectory, cells acquire GABA-OPC hybrid identity through pseudotemporal gains of *GAD1*, *PDGFRA* and *PTPRZ1*, consistent with the capacity of astrocytic progenitors to generate OPC-like daughter populations through instructive fate decisions ^66^. Along the second trajectory, cells acquire markers of neuronal-committed NPC biology (*DCX*, *TUBB3*, *SOX11*), consistent with post-mitotic neuronal commitment ^67–69^. This developmental logit parallels radial glial lineage biology, where astrocyte-like radial glial cells generate transcriptionally diverse progenitor populations ^60^. The third and fourth trajectories are the most clinically consequential, culminating in Mitotic Proliferative and Mesenchymal endpoints, respectively. The third putatively proceeds through a Proteo-Synthetic intermediate, with pseudotemporal gains of key markers (*CENPE*, *KIF11*, *MKI67*) toward the terminus ^70–72^. The final trajectory proceeds toward the Mesenchymal endpoint through acquisition of TGFβ-driven motility programs, NCOR2-mediated chromatin remodeling, and MEF2-family regulatory control ^73–75^.

To strengthen our pseudotime dynamic claims across disease progression, scaled values were assessed in four patients with matched primary and recurrent specimens (**Supplementary Figure S4a-d**). In three of four patients, scaled pseudotime increased monotonically from primary GCH through recurrent GCL timepoints. State abundance shifts at GCL recurrence showed contracted Astrocytic-like representation and expanded terminal state abundance concordant with trajectory predictions across all patients. While the small number of longitudinal cases precludes definitive conclusions, these observations support the multifurcating developmental architecture as a reproducible, patient-intrinsic feature of G-CIMP disease progression.

### Cellular state assignment is an independent predictor of overall survival

To determine the relation of our cellular states to clinical outcome, multi-center bulk transcriptomes from IDH-mutant glioma (**Table 1**; TCGA, CGGA, and GLASS; N=478) were classified according to predominant metaprogram (**Supplementary Figure S5a-c**). Kaplan-Meier survival curves saw significantly poorer survival for Mitotic Proliferative and Mesenchymal samples (log-rank p<0.0001; **Figure 4a**). These states retained independent prognostic significance across a multivariable Cox proportional hazards model, adjusting for WHO grade and age [Mitotic Proliferator: HR=2.53, 95% CI: [1.54-4.14], p<0.001; Mesenchymal: HR=4.25, 95% CI: [2.34-7.74], p<0.001)(**Table 2**, **Supplementary File S3**). Additional adjustment for MGMT-promoter methylation status across samples with available data (n=327) had no impact. Distribution of clinicopathological features across states reinforced nomenclature validity; higher mortality in Mitotic and Mesenchymal states, Astrocytic-like state concentration of low-grade astrocytomas, oligodendroglioma saturation within GABA-OPC samples, and predominance of grade 4 within the Mitotic group. The Mesenchymal state was strikingly exclusively composed of IDH-mutant non-codeleted tumors, consistent with the known association between codeletion and mesenchymal transcriptional programs ^4^.

**Figure 4.**
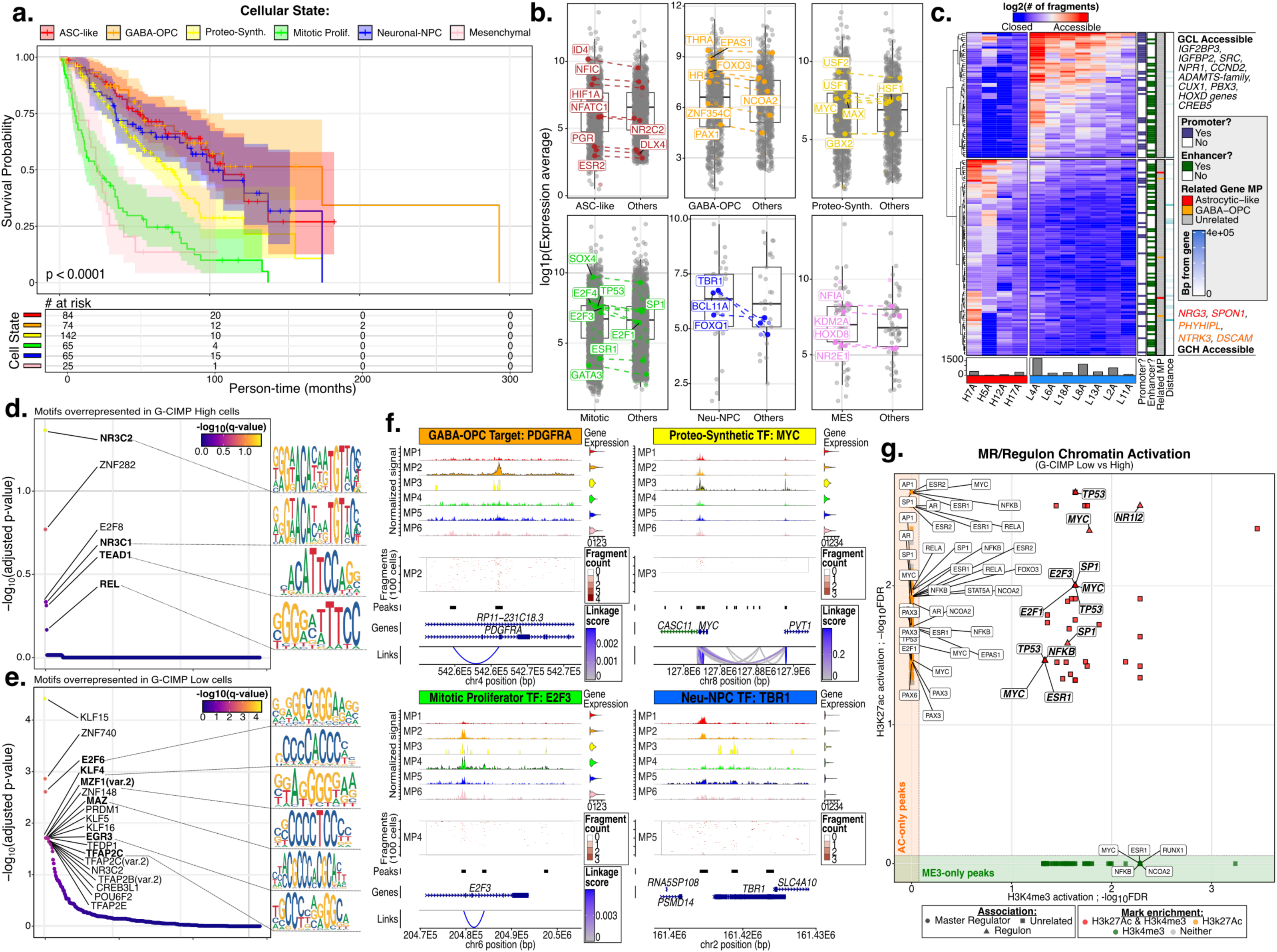
Exploration of snRNA-seq cellular states across additional omic data types (bulk RNA-seq, snATAC-seq, ChIP-seq). **a)** Multi-center Kaplan-Meier survival curves stratified by assigned cellular state, depicted with 95% confidence intervals (vertical ticks: censorship). **b)** Boxplots displaying the expression of inferred master regulators (MRs) and putative regulons, across bulk samples. **c)** Heatmap displaying snATAC-seq differential peaks. Columns are denoted by the number of cells, and accessible genes are displayed. **d-e).** Ranked order plots displaying estimated significance of over-represented motifs discovered through pseudobulking across **(d)** G-CIMP-high and **(e)** G-CIMP-low cells. **(f)** Coverage plots of differential peaks related to cellular states. **g)** Scatterplot depicting the activation of cellular state MR and associated regulons across H3k27Ac and H3k4Me3 datasets. Points are shaped according to gene type and colored according to significance.

**Table 1.** Clinicopathological feature distributions of multi-center bulk IDHmutant glioma stratified by cellular state.

**Table 2.** Multivariable cox proportional hazards model - overall survival.

### External bulk data sources provide multi-modal validation of snRNA-seq transcriptomic findings

To validate the transcriptional activity of proposed MRs, we examined their expression across CGGA IDH-mutant glioma bulk transcriptomes (n=356), stratified by cellular state ^76^. Expression was heightened in affiliated states of origin, supporting state-specific transcriptional activity of these regulators (**Figure 4b**).

Additional external multi-omic validation confirmed state-specific clinicopathological feature concentrations ^77^. For example, WHO Grade 4 tumors were predominantly associated with the Mitotic Proliferator state. In paired scRNA-seq data, bulk-derived cellular state assignments showed limited concordance with state abundance, with the exception of one sample showing Mitotic Proliferator dominance, reflecting bulk transcriptomics inherent limitations (sample: CGGA_P1088.2; **Supplementary Figure S5d**). Across states with differential distributions of both MP gene expression and associated protein abundance (n=4 of 5), significant correlations between both modalities were observed (Spearman ρ range: [0.32-0.4], p<2.2×10^-16^; **Supplementary Figure S5e**). Widespread promoter-region CpG hypomethylation was revealed for three cellular states (Astrocytic-like, Proteo-Synthetic, Mitotic Proliferator). Together these modalities converge in support of a coordinated multi-omic regulatory architecture across associated transcriptional programs in independent IDH-mutant data sources.

Furthermore, we observed notable coordinated gene expression architectures arising from particular MRs and their putative target genes within bulk transcriptomics. For example, Neuronal-NPC MR *TBR1* showed confluent upregulation of predicted target genes (>50%) and regulator strength in bulk sources (**Supplementary Figure S5f**).

### Chromatin accessibility and histone modification converge on GCL regulatory hubs

Following isolation of acceptable-quality nuclei across the single-nucleus accessibility profiling (snATAC-seq) data set and pseudobulking to overcome data sparsity, differential accessibility analysis identified regions of increased chromatin accessibility in GCL proximal to genes associated with proliferation (*IGF2BP3*, *IGFBP2*, *CCND2*), angiogenesis (*NRP1*) and therapy resistance (*SRC*, *RAD51*), among others (**Figure 4c**, **Supplementary Figure 5g**) ^78–82^. These findings are consistent with the transcriptional programs identified in GCL-specific cellular states. Conversely, GCH samples showed increased accessibility at Astrocytic-like (*NRG3*, *SPON1*) and GABA-OPC (*PHYHIPL*, *NTRK3*, *DSCAM*) markers, consistent with the enrichments of less aggressive states (**Figure 4c**).

Motif enrichment analyses of GCH-enriched regions possessed motifs corresponding to nuclear receptors and stress-responsive TFs including *NR3C1/2* and *REL*, consistent with chromatin landscapes configured to favor differentiation and stress signaling programs (**Figure 4d**) ^83^. Conversely, GCL-enriched accessible regions identified TFs implicated in proliferation, cell-cycle regulation and lineage specification (e.g., E2F-family members, *KLF4*, *MAZ*) (**Figure 4e**) ^84,85^. State-specific snATAC-seq accessibility corroborated MR predictions: Proteo-Synthetic cells showed increased *MYC* locus accessibility, the GABA-OPC state saw *PDGFRA* locus enrichment, and Mitotic Proliferators recapitulated E2F-family accessibility (**Figure 4f**).

Bulk H3K27ac/H3K4me3 ChIP-sequencing across independent tumor cohorts confirmed dual-mark activation at *MYC*, *E2F1/3*, *NHLH2* and HOXD-family loci in GCL tumors, providing multi-modal epigenomic validation of the transcriptional regulatory architecture identified through snRNA-seq (**Figure 4g**, **Supplementary Figure 5h**). Particularly, simultaneous concordance of snATAC-seq accessibility enrichment at *NHLH2* and *HOXD13* with ChIP-seq signal across an independent cohort provides cross-platform convergence for GCL-specific regulatory activation at these loci, representing tractable starting points for functional investigation of cis-regulatory mechanisms underlying the G-CIMP epigenomic transition.

### Immune compartment stratification and characterization

Following immune component identification, we sought to functionally stratify immune cell populations within the tumor microenvironment (**Supplementary Figure S6a)**.

Cross-atlas program enrichment analysis using curated immune signatures derived from published single-cell studies was performed, and shared enrichments for reference programs interpreted as higher-order identities, yielding six “super clusters” (**Figure 5a, Supplementary Figure S6b**) ^86–88^. Additionally, we sought to functionally characterize the clusters utilizing pseudobulked expression datasets (**Figure 5b-c**, **Supplementary Figure S6c**).

**Figure 5.**
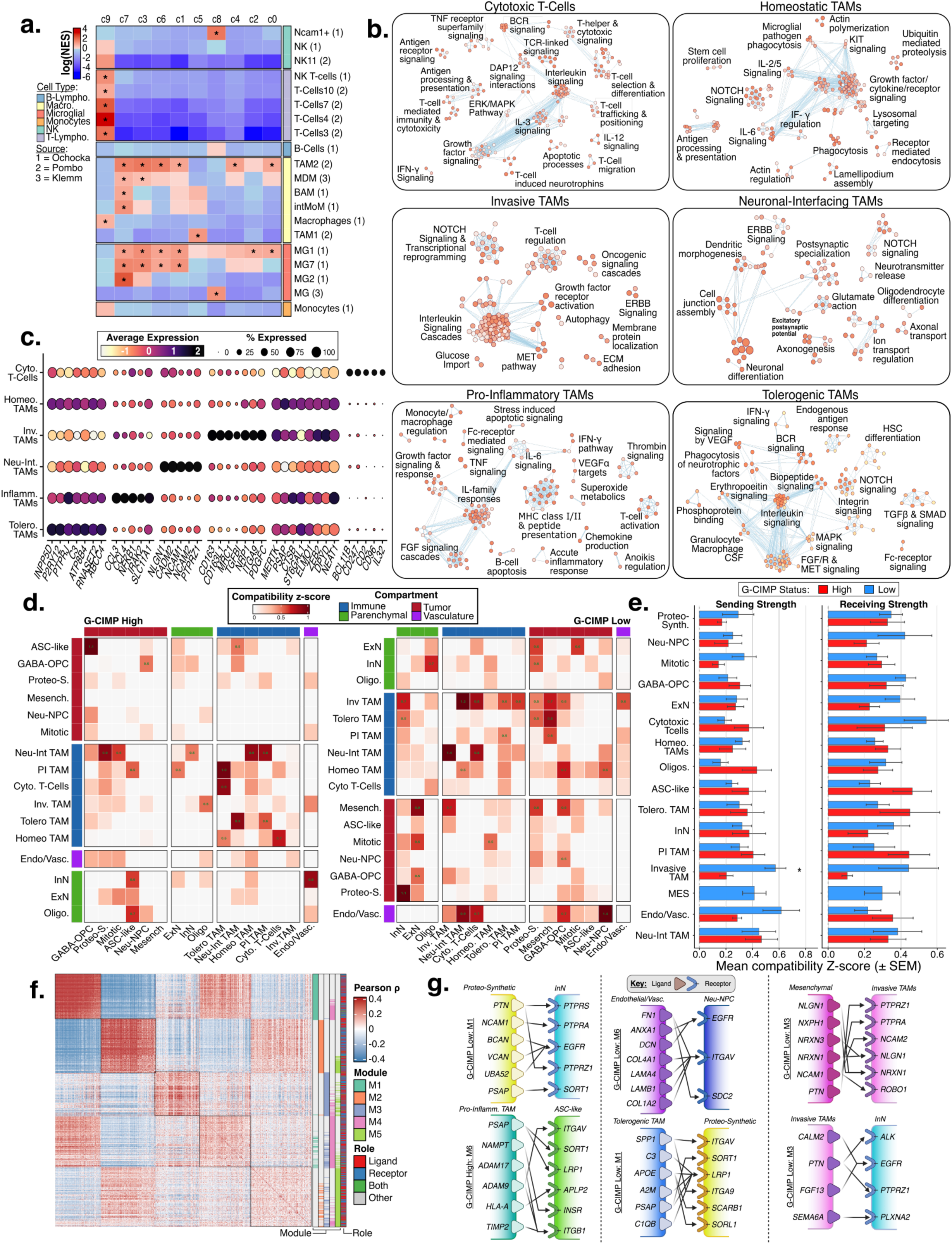
Definition of the Immune tumor-microenvironment and inferred communication with malignant states. **a)** Heatmap depicting reference immune cell type transcriptional signature enrichment across clusters. **b)** EnrichmentMap networks for the identified immune super-clusters. **c)** Dot plot displaying relevant marker expression across cells, grouped by immune supercluster. Dots are colored by the average expression value, and sized by the percentage of cells expressing each marker. **d)** Heatmap series displaying inferred compatibility z-score values between sending cells (y-axis) and receiving cells (x-axis), split by G-CIMP. **e)** Barplot series depicting mean values of compatibility scores attributed to total sending and receiving activity across cell populations, stratified by G-CIMP. Note: SEM: Standard Error of the Mean. **f)** Correlation based heatmap displaying modules of correlated ligand and receptor molecules through interaction between Proteo-Synthetic and InN populations in G-CIMP-low tumors. Molecules are annotated by module membership and signaling role. **g)** Schematic displaying modules of ligand and receptor interactions across specific cell type combinations.

The first supercluster, designated ‘tolerogenic’ tumor associated macrophages (TAMs), possessed immune tolerance and environmental adaptation ontologies, alongside microglia-associated markers (*P2RY12*, *ATP8B4*) and complement-associated immune modulation (*C3*) ^88,89^. This regulatory and anti-inflammatory identity was the sole supercluster displaying differential abundance across clinicopathological features, preferentially enriched in GCH and low-grade tumors (**Supplementary Figure S6g**). The second, namely homeostatic TAMs, was defined by environmental surveillance programs, lysosomal pathways, Fcγ receptor signaling and a *ZEB2*-mediated regulatory identity ^90,91^. The third super-cluster, namely ‘invasive’ TAMs, displayed activation of matrix interaction programs, suggested responsiveness to tumor-derived growth factors and expressed ECM-interacting molecules (*TGFBI*, *ITGA9*) and angiogenic receptors (*NRP1*), supporting a tissue-invasive classification ^92,93^. The fourth population, ‘neuronal-interfacing’ TAMs, expressed synaptic adhesion programs and *PTPRZ1*, demonstrating transcriptional acquisition of neuro-active programs inconsistent with stochastic neural contamination and consistent with emerging evidence for macrophage participation in tumor-neural interfaces ^94,95^. The fifth, designated pro-inflammatory TAMs, were distinguished through simultaneous activation of classical inflammatory pathways and cytokine-mediated communication, alongside expressed inflammatory cytokines (*NFKB1*, *IRAK2)* and oxidative stress regulator *SCL11A1* ^96,97^. Our final population, a T-cell supercluster, was distinguished by presence of immune-mediated cytotoxicity and expression of markers consistent with classical T-cell identity (*BCL11B*, *IL32*) and functional activation (*CD247*, *CD96*) ^98,99^.

### Immune microenvironment reorganization accompanies G-CIMP epigenomic transition

To characterize intercellular signaling across G-CIMP strata, we applied a custom two-stage ligand-receptor compatibility and module discovery pipeline. Compatibility z-scores, defining the degree to which sender-ligand and receiver-receptor expression profiles are coordinately expressed, revealed an expanded predicted communication architecture in GCL tumors relative to GCH. (**Figure 5d-e**).

Summarily, GCL tumors displayed high compatibility across immune-tumor and tumor-parenchymal axes. Invasive TAMs emerged as prominent senders and receivers, consistent with prior detailed enrichment of pro-tumorigenic immune populations in high-grade IDH-mutant contexts ^100,101^. Enhanced outgoing compatibility in the endothelial/vascular compartment was consistent with a vascular niche construction axis ^102,103^. Parenchymal GCL neuronal populations (ExN, InN) showed preferential receipt when compared with GCH, suggesting the surrounding neural parenchyma as more broadly engaged, consistent with prior reports ^104,105^.

For each candidate sender-receiver population pair identified through pre-screening, detailed co-expression module identification was completed to characterize internal gene program architecture (**Figure 5f-g; Supplementary File S4**). Key predicted axes included tumor-neural interface remodeling via *BCAN*/*VCAN*-*PTPRS1* and *PSAP*-*SORT1* interactions in a Proteo-Synthetic to InN axis, and vasculature to tumor (Neuronal-NPC) signaling denoting perivascular niche construction through interactions of *ITGAV*-*EGFR* and vascular secretion of *ANXA1* ^106–112^. Notable Mesenchymal tumor signal received by invasive macrophages saw an outgoing trans-synaptic adhesion complex (*NLGN1*, *NXPH1*, *NRXN1/3*), suggesting a candidate signaling axis through which mesenchymal tumor cells interface with pro-invasion TAMs ^113,114^. GCH context pro-inflammatory macrophage signal to Astrocytic-like tumor cells similarly saw directional interfacing, through which lysosomal enzyme delivery and biosynthetic support signaling were achieved through interactions such as *PSAP*-*SORT1* and *NAMPT*-*LRP1* ^115–118^. Potential metabolic provisioning and support through *SPP1*-*ITGAV* and downstream effectors (*A2M*-*LRP1*, *SCARB1*) was discovered between tolerogenic TAM and Proteo-Synthetic GCL populations ^119–123^. GCL Invasive macrophages presented unconventionally secreted extracellular factors in *CALM2*, activating *ALK* on inhibitory neurons, alongside growth factor (*EGFR*) activation through *PTN*/*FGF13*, and simultaneous PTPRZ1 inhibition; altogether, interactions were consistent with additional remodeling of the tumor-neural interface ^124–128^

Inquiry into the ontological background present across the discovered modules was conducted through compartment stratified GO specificity analyses (**Supplementary File S4**). Briefly, analysis confirmed interaction-specific biological thematics across immune-tumor, tumor-parenchymal and vasculature-tumor compartments, consistent with the glioma microenvironment. For example, Immune-to-tumor interactions were specifically enriched for neuroinflammatory response, regulation of adaptive immune response and leukocyte activation processes, consistent with TAM-mediated immunomodulatory signaling in the glioma microenvironment ^101,129^. Additional tumor-to-parenchymal interactions were dominated by *ERBB2*/*ERBB4* signaling, transmembrane receptor tyrosine kinase activator activity and synaptic-membrane related terminology, reflecting the shared neural transcriptional identity of tumor and parenchymal populations and summarizing module-level findings.

## Discussion

This study provides a single-cell resolution portrait of the G-CIMP epigenomic transition in IDH-mutant glioma, resolving six recurrent malignant states, their chromatin accessibility coordinates, a multifurcating developmental model, and a fundamentally reorganized tumor microenvironment between GCH and GCL contexts. These findings provide mechanistic grounding for GCL-associated poor prognosis, and illuminate molecularly tractable regulatory nodes for therapeutic interception.

NMF-based decomposition of the malignant transcriptome yielded six robust metaprograms whose differential enrichment across G-CIMP strata and WHO Grades recapitulated and extended prior single-cell IDH-mutant and IDH-wildtype glioma studies ^12,20,39^. Dominance of the Astrocytic-like state in GCH tumors was consistent with prior observations that GCH tumors maintain differentiated cellular programs analogous to their presumed cell of origin ^6,10,130^. Conversely, the differential enrichment of Mitotic-Proliferative and Mesenchymal states in GCL tumors aligns with the epigenomic erosion model: progressive demethylation and enhancer remodeling drive lineage infidelity and re-acquisition of stem-like plasticity ^8,131^. Within this framework, the GCL epigenome does not simply “lose” the GCH signature, but actively acquires a permissive regulatory landscape, most consequentially enabling the Mesenchymal state, representing a GCL-specific cellular identity absent from prior IDH-mutant reference taxonomies, exclusively non-codeleted in external validation, and carrying the worst prognosis. Direct pseudotemporal trajectory from the Astrocytic-origin may reflect selective *MEF2*/TGFβ enhancer erosion at the GCL transition.

The positioning of the Proteo-Synthetic state pseudotemporally as an intermediate preceding mitotic commitment, arranges biosynthetic groundwork as a prerequisite for proliferative expansion in GCL tumors ^132^. Co-option of this program, potentially *MYC*-driven, positions this state as a candidate transitional stage whose disruption may prevent downstream Mitotic Proliferative commitment. Additionally, the GABA-OPC hybrid state likely represents a stable lateral differentiation endpoint accessible regardless of epigenomic subtype; sustained by a *FOXO3*/*CEBPB* quiescent progenitor program, broad distribution and clinically intermediate prognosis warrants investigation in larger cohorts.

The delineation of a putative developmental model reveals that the G-CIMP transition corresponds to a probabilistic redistribution of trajectory endpoint utilization, where GCL tumors gain disproportionate access to proliferative and mesenchymal termini at the expense of quiescent and differentiated branches ^4,6^. While the small number of validative longitudinal cases precludes definitive conclusions, this model provides a candidate pharmacodynamic readout against which the developmental impact of vorasidenib could be prospectively quantified through longitudinal biopsy state distribution analysis.

Convergence of master regulator inference, snATAC-seq motif enrichments, and bulk ChIP-seq across independent cohorts onto *E2F*, *MYC*, *MEF2* and NOTCH regulatory hubs defines a well supported therapeutic landscape ^133^. Potential *CDK4/6* inhibition targeting the E2F apex, particularly through brain-penetrant agents such as ademaciclib, represents the most immediately translatable intervention ^134,135^. *KIF11*, whose expression correlates with WHO grade and whose inhibition by ipinesib reduces glioblastoma proliferation and invasion in orthotopic models, is pseudotemporally co-gained with *MYC*-stabilizing protein *CIP2A* at the proliferative terminus, defining a transition-point vulnerability ^72,136^. *BET* bromodomain inhibition simultaneously targets biosynthetic priming and downstream E2F-driven enhancer programs ^137^. NOTCH pathway interference through gamma-secretase inhibitors or *CBF1*/*RBPJ* blockade, motivated by independent co-gain of Mesenchymal terminus effectors *MAML2* and *RBPJ*, addresses the Mesenchymal adaptation specifically ^138,139^.

Here, limitations include: modest cohort size (n=10 patients, 18 specimens), inability to fully decouple G-CIMP status from potentially confounding factors such as WHO grade and treatment history, below-standard snATAC-seq transcription start site (TSS) enrichment, and inferential nature of predicted ligand-receptor interactions and developmental trajectories, which require functional confirmation through lineage-tracing and co-culture approaches ^140–142^. Future work with larger histology-stratified and treatment-naïve cohorts should prioritize spatial transcriptomics, longitudinal single-cell profiling through IDH-inhibitor trials, and functional validations of inferred regulatory, developmental, and microenvironmental programs in patient-derived models.

## Conclusion

This study provides a multimodal, single-nucleus resolution map of the malignant and microenvironmental cellular ecosystems which distinguish G-CIMP-low from G-CIMP-high in IDH-mutant gliomas. We identify six recurrent malignant cellular states which exhibit differential G-CIMP enrichment, convergent chromatin accessibility and histone modification signatures, and distinct pseudotemporal roles within a multifurcating developmental model anchored in a GCH Astrocytic-like origin. Mesenchymal and Mitotic Proliferative states are independently prognostic in external cohorts and define the clinically consequential terminal endpoints of GCL trajectory redistribution. Six immune superclusters, including a novel neuronal-interfacing macrophage population, and an expanded predicted communication matrix in GCL tumors reveal that the epigenomic transition is accompanied by co-evolution of a permissive multi-compartment microenvironment. Cross-platform regulatory convergence on E2F, *MYC*, *MEF2*, *KIF11* and NOTCH axes provide an immediately actionable therapeutic framework. These findings advance IDH-mutant glioma single-cell characterization from descriptive taxonomy toward a mechanistically grounded, developmentally anchored, and translationally oriented framework for intercepting GCL progression in this clinically underserved population.

## Methods

### Human Subjects

This project was approved by the Institutional Review Boards (IRB) and patients consented to have their specimens used for research purposes (IRB #15367). We profiled specimens who underwent resection at the Neurological Surgery Department at Henry Ford Health from 08/1994 to 01/2020 (n=18 samples). Longitudinal collections were available for four patients, with two patients contributing primary, first-recurrent and second-recurrent samples. Profiling was completed in two separate efforts/batches, the first consisting of eight and the second consisting of ten collections completed on 05/2023 and 06/2023, respectively. The second effort contained two replicated samples (namely: L11C & L13C), for quality assurance purposes. Specimens collected from IDH-mutant patients were previously stratified by glioma CpG island methylator phenotype or ‘G-CIMP’ status through bulk methylation profiles ^143^. The cohort consists of thirteen high-grade gliomas (HGG; anaplastic astrocytoma: n=1, glioblastoma: n=7, anaplastic oligoastrocytoma: n=2, anaplastic oligodendroglioma: n=3) and five low-grade glioma (LGG; astrocytoma: n=1, oligoastrocytoma: n=2, oligodendroglioma: n=2). Nine samples were resected from primary tumors and nine from recurrent tumors (first recurrence: n=4; second recurrence: n=5). Four samples were collected from patients diagnosed between the ages of 18-30, fourteen from patients aged 30-50 years old, and four samples from patients older than 50. Complete clinical information is available in **Supplementary File S1**.

### Nuclei Isolation from Frozen Brain Tumor Tissue

Nuclei were isolated from 10mg of frozen brain tumor tissue using the *10x Genomics Nuclei Isolation for Single Cell Multiome ATAC + Gene Expression* demonstrated protocol (Document CG000365, Rev D), with minor modifications. All buffers were prepared fresh and maintained at 4°C; full buffer compositions and procedural details are provided in **Supplementary Methods**.

### Single-Cell Multiome ATAC + Gene Expression

Single-cell 10x Genomics Multiome (ATAC + gene expression) libraries were generated using the 10x Genomics Chromium Single Cell Multiome platform from human nuclei (hg38), targeting ∼4,000 nuclei per sample. Libraries were sequenced by Psomagen (Illumina NovaSeq 6000, S1 v1.5 flow cell) at 10x-recommended read lengths (28 × 10 × 10 × 90). Ten libraries were pooled at equimolar ratios and sequenced on a single flow cell. FASTQ files were generated using cellranger-arc mkfastq. Specific details can be found in **Supplementary Methods**.

### snRNA-seq Data Processing

Raw sequencing reads were aligned to the GRCh38 human reference genome (Ensembl release 2020-A) using Cell Ranger v7.0.1 (10x Genomics). Processing and quality control filtering of the snRNA-sequencing dataset was conducted as previously described, with some experiment-specific alterations ^144^. Briefly, cells with fewer than 1,000 detected genes were filtered out from each sample, resulting in a median of 1,850 detected genes per cell across the dataset, including both neoplastic and non-neoplastic populations. To exclude low-quality or dying cells, we filtered based on the percentage of reads mapping to the mitochondrial genome (<3%; GQ cell median %: 0.248), and ribosomal proteins (<5%; GQ cell median %: 0.707). These filtration steps resulted in a median cell count per sample of 947.5 cells (Q1: 620.75, Q3: 1752.5), with a single sample retaining less than 500 total cells (H12).

Cells surpassing quality control thresholds were clustered following initial dimension reduction (Principal Component Analysis; PCA) on scaled expression values for the 2,000 most variable genes. Principal components were ranked by the percentage of variance explained, and the top components were retained for downstream analysis. A K-nearest neighbors graph was constructed based on Euclidean distances in PCA space, and cells were clustered using the Louvain algorithm as implemented in Seurat v5.3.0 (*FindClusters*; resolution=0.25) ^19^. Clusters were visualized in two dimensions using Uniform Manifold Approximation and Projection, or UMAP, computed on the same principal components.

### Neoplastic and non-neoplastic cell discrimination and cell type annotation

Following identification of clusters for each sample, initial exploration of cell phenotype was conducted using the scType framework ^145^. Briefly, using curated signature sets for selected CNS cell subtypes, each cell was assigned an enrichment score based on the expression of both positive and negative markers. Then, each cluster was designated a consensus label based on the frequency of assigned phenotypes. Clusters in which fewer than 25% of cells shared a common phenotype were labeled ‘Unknown’ and excluded from annotation. Classifications of oligodendrocytes, immune system cells, microglial cells, endothelial cells, GABAergic and glutamatergic neurons were considered potential non-neoplastic phenotypes. Neoplastic cell types included radial glial cells, oligodendrocyte precursor cells (OPC), astrocytes, neuroblasts, and select neuronal populations (mature/GABAergic).

To infer copy number alterations (CNA), we used inferCNV (v1.24.0) across gene expression profiles per sample, using non-tumoral clusters as the reference population, with gene positions defined through GRch38 Ensembl annotation ^146^. The algorithm sorts genes by chromosomal location and computes a moving average of relative expression values through utilization of a 100 gene-width sliding window. Additionally, *absolute* CNA values were computed using inferCNA (v1.0.0), which defined two separate scores: first, ‘correlation’ which defines the Pearson correlation between the CNA profile of each individual cell with that of the average CNA profile of the sample’s tumor cells; second, ‘signal’, defined as the sum of CNA values across all genes for each cell. Both measures were utilized for each individual tumor to segregate neoplastic from non-neoplastic cell types, for example those with high correlation and signal being treated as neoplastic cells, and those with the inverse being treated as non-neoplastic. Any discordant cells were excluded from downstream analyses (n=5,999 cells).

### Definition of neoplastic cell metaprograms

Low-variance ribosomal protein genes (*RPL*- and *RPS*-family members) were excluded prior to Non-negative Matrix Factorization (NMF) decomposition, as their high constitutive expression reflects global translational activity rather than biologically informative transcriptional variation. NMF was employed to identify and define neoplastic cell MPs following the framework established in Gavish et al., 2023 [Citation error]. In this approach, NMF decomposes gene expression matrices into sets of co-expressed genes (metaprograms) and their corresponding cell-level activity scores, capturing recurrent transcriptional states across tumor cells. Summarily, NMFs were applied to the log-normalized expression values of each sample, with any negative expression values set to zero. Iterations were completed for every value of expected number of clusters across each sample between k=4 to 9, generating 39 distinct NMF programs for each sample (n=702 total). Each was then defined based on the 100 genes which possessed the highest estimated NMF coefficients. We then retained only those programs which could be defined as robust within and across samples, and non-redundant within the sample, as previously defined [Citation error].

The resulting number of robust and non-redundant NMF programs was 131, which were clustered according to Jaccard similarity using the degree of overlap between programs, ultimately combining those which possessed high degrees of overlap. MP signatures were then defined as the 100 most recurring genes across the programs within a shared Jaccard cluster, leading to a final number of six robust and non-redundant biological metaprograms. Although MP3 showed relatively high ribosomal gene contributions, it satisfied all implemented robustness and non-redundancy criteria and was retained as a putative biological program.

### Classification of neoplastic cells into metaprograms

Neoplastic cells were defined as belonging to one of our identified metaprograms using a combinatory approach, adapted from previous efforts [Citation error]. First, a non-parametric one-sided Wilcoxon rank-sum test was completed for each cell across each MP gene list, comparing the expression of the selected genes to background activity. Derived p-values were then adjusted for false-discovery rate (FDR). Then, we applied Seurat’s *AddModuleScore*, with scores assigned to each cell for each MP ^19^. Cells were designated according to MP through a combination of maximum module score, and a significance threshold of FDR-adjusted p≤0.05. If a cell did not possess significance at the maximal module score, it was left ‘Unassigned’.

### Characterization of cell metaprograms and cell cycle phases

To further characterize the identified cellular states, we explored functional enrichments, correlations with published transcriptional programs, and cell cycle assignments.

Functional enrichment testing was performed using gProfiler2 across *Homo Sapiens* collections (GO:BP/MF, KEGG, Reactome, WikiPathways; gene set size >3, intersection size >5, FDR<0.05) using the transcriptional MPs (n=100 genes, **Supplementary File S2**) ^147^. Enriched terms were matched to the Bader Labs Human GO collection and filtered to retain brain, tumor, and immune relevant processes ^148^. In parallel, rank-based gene set enrichment analysis (GSEA) was performed using yaGST (v2017.08.25; https://github.com/miccec/yaGST) to derive normalized enrichment scores (NES) as a complementary measure of effect size. Functional similarity networks were constructed in Cytoscape’s EnrichmentMap (FDR<0.05, Jaccard>0.25) and annotated through AutoAnnotate ^149^.

Two complementary correlation approaches were applied across all malignant cell populations. For comparisons with normal brain cell type programs, module scores for reference cell type signatures were computed using Seurat’s *AddModuleScore*, drawn from six published normal brain single-cell atlases [Citation error]. Per-cell module scores were correlated with metaprogram module scores (MP1-MP6) using Pearson correlation; normal brain signatures with a maximum absolute correlation exceeding 0.6 with any metaprogram were retained for visualization. For comparisons with previously published malignant cell states, three single-cell glioma reference atlases were used [Citation error]. For each reference atlas, one-sided Mann-Whitney Wilcoxon enrichment scores (logNES, package: yaGST) were computed per cell per reference program. Cells were assigned to the reference program with the maximum logNES (p≤0.05), or left ‘Unassigned’. Pairwise Jaccard similarity indices were computed between our six metaprograms and each set of reference assignments using the intersection over union.

Cell cycle phase assignment was completed using ccAFv2, a program which allows for the assignment of six phases: G1, Late G1, S, S/G2, G2/M, and M/Early G1, and a quiescent-like G0 state (Neural G0) [Citation error]. In attempts to draw informative conclusions, we consolidated the phases into four primary categories: G1 (G1, Late G1), S (S, S/G2), M (G2/M, M/Early G1), and Neural G0. Those cells which did not fall into any phase with high confidence (score<0.5) were labeled as ‘Unknown’. We further consolidated cells into a tertiary system of ‘Active’ (S-, M-phase), ‘Inactive’ (G1-phase) and ‘Quiescent’ (Neural G0) to facilitate interpretation of proliferative versus quiescent cellular states within the tumor microenvironment, given small per-phase cell counts in some metaprograms.

### Discrimination and characterization of neuronal-related malignant cellular states

Due to overlapping correlations of MP2 and MP5 cells with neuronal reference programs, a series of complimentary analyses were undertaken to resolve their distinct identities.

First, differential expression testing between cells was performed using Seurat’s *FindMarkers [Citation error]*. Resulting genes were ranked by average log_2_(fold-change) and annotated to neurotransmitter receptor families based on established membership (**Supplementary File S2**). To systematically assess correlations with established neuronal subtypes, a master reference set was compiled from published atlases ^150^. For each cell, Pearson correlations were computed between the average log-normalized expression of each reference signature and a binary metaprogram membership vector. Reference signatures were classified as inhibitory, excitatory, or unknown based on identifier keywords. Correlation rank plots with the top five signatures labeled were generated per metaprogram. Correlation values were then compared between MP2 and MP5 across excitatory and inhibitory signature sets using Wilcoxon rank-sum testing, and visualized through boxplots.

To further separate MP2 and MP5 using a multi-program scoring approach, one-sided Mann-Whitney Wilcoxon enrichment scores (logNES; package: yaGST) were computed per cell for OPC, InN and ExN reference programs (**Supplementary File S2**) [Citation error].The resulting score matrix was subjected to PCA, and cells were visualized across components and colored by metaprogram identity, OPC, ExN and InN scores, respectively.

To confirm the hybrid oligodendrocyte-GABAergic identity of MP2 cells, a co-expression analysis was performed across MP2 malignant cells, non-malignant InN and oligodendrocytic populations. Non-malignant population-specific markers were identified through Seurat’s *FindAllMarkers* ^19^. Identified signatures were visualized through an expression based heatmap. Additional marker-level validation was completed using curated gene lists (**Supplementary File S2**). Finally, enrichment scores for three signature sets: (i) a curated GABA-OPC signature, (ii) a normal OPC signature and (iii) a consensus cross-atlas oligodendrocytic signature (genes present in ≥2 atlases across sources) were computed for each MP2 cell, and enrichments were compared by Wilcoxon rank-sum test [Citation error].

To characterize pathway-level differences distinguishing MP2 malignant cells from non-malignant oligodendrocytes, and MP5 malignant cells from non-malignant neurons, separate differential expression analyses were performed for each comparison (Seurat’s *FindAllMarkers*) ^19^. Significantly upregulated genes in each malignant population relative to its non-malignant counterpart were submitted to Gene Ontology enrichment analysis using clusterProfiler’s *enrichGO* function ^151^.

To confirm the neuronal-committed progenitor identity of MP5, comparative analysis was performed against the mitotic-proliferative state (MP4). This comparison was selected on the basis that MP4 cells represent the most proliferatively active progenitor-like state within the malignant compartment. Three custom canonical progenitor marker sets were curated, namely: (i) neuronal-committed NPCs, (ii) neural stem cells and (iii) multipotent/uncommitted NPCs (**Supplementary File S2**). Module scores (Seurat’s *AddModuleScore*) and expression plots were generated per program, with statistical comparisons of module scores performed using two-sided Wilcoxon rank sum testing ^19^.

### Trajectory Analysis, Batch Correction and Pseudotime Inference (Slingshot)

In efforts to investigate the underlying developmental lineage of our malignant cell cohort, particularly through the lens of G-CIMP state transition, we implemented Slingshot (v2.16) ^63^. Briefly, a subset of tumor samples were selected, following the exclusion of samples with fewer than 250 malignant cells (final cohort: n=12 samples). Batch effects arising from inter-sample technical variations were corrected using Harmony (v1.2.4), and the resulting corrected embedding was utilized for all downstream dimensionality reduction, graph construction and trajectory inference ^64^.

Pseudotime trajectory inference was performed operating on the Harmony-corrected PCA embedding (n=15 dimensions), to avoid the non-linear topological distortions introduced through UMAP. The Astrocytic-like state was identified as the fixed trajectory origin, based on its preferential enrichment in GCH primary tumors, quiescent G0/G1 cell cycle profile, and consistency with established glioma developmental hierarchies ^12,152^. Four terminal states were specified a priori based on biological rationale and G-CIMP-stratified abundance analyses: namely: GABA-OPC, Mitotic Proliferator, Neuronal-committed NPC, and Mesenchymal cell states.

A unified global pseudotime variable was derived for visualizations using the Slingshot-native weighted average pseudotime. Resulting values were linearly rescaled to the unit interval [0,1] to facilitate cross-sample comparison and visualization. Cells not assigned to any lineage branch (curve weight<0.1) were excluded from pseudotime visualizations.

### Pseudotemporally Dynamic Gene Identification

To identify genes with significant expression dynamics across each of the identified lineages, branch-specific generalized additive models (GAMs) were fitted using tradeSeq (v1.22.0) ^65^. For each branch, cells were selected based on curve weights (>0.3), isolating branch-specific cells.

Gene pre-selection for each branch was performed in two stages. First, genes were retained if expressed in a minimum fraction of terminal-state cells (GABA-OPC: 15%, Mitotic/Neuronal-NPC: 20%, Mesenchymal:50%). Expression thresholds were branch specific to account for differences in cell composition and terminal state representation. Second, branch-specific highly variable genes were identified through scran’s function *modelGeneVar*, applied to branch cells, with top-ranked genes selected (DOI: <u>10.18129/B9.bioc.scran</u>). Genes matching patterns indicative of technical or uninformative biological signal were excluded prior to all analyses, including ribosomal protein subunits, histone genes and variants, transcription elongation and initiation factors, ubiquitin pathway components, and common housekeeping genes (*GAPDH*, *ACTB*, *B2M*, *FTL*).

Terminal marker identification was performed, and determined whether expression was gained or lost toward each developmental culmination point, using tradeSeq’s *startVsEndTest*. We also complementarily applied *associationTest* to capture intermediate dynamic genes. Given the exploratory nature of terminal marker identification and the prior applied gene filtering, an unadjusted p-value threshold of 0.05 was used. Results were interpreted in conjunction with Wald statistic magnitude and prior biological knowledge rather than as a definitive significance screen. For visualizations, curated marker genes were selected per branch based on a combination of statistical strength (Wald statistic), biological relevance to the terminal state identity and specificity relative to the origin state.

### Longitudinal Pseudotime Validation Across Matched Primary and Recurrent Samples

To validate pseudotime dynamics across disease progression, values were assessed in patients with matched primary and recurrent specimens (n=4). For samples not included in the pooled trajectory model, pseudotime was estimated via reference-based projection and inverse-distance-weighted k-NN interpolation (k=15) onto the pooled Harmony embedding; full projection details are provided in **Supplementary Methods**.

For visualization of pseudotime measures, inter-timepoint batch effects were corrected through Harmony. UMAP dimensionality reduction was computed on the patient-specific Harmony-corrected embedding. Cellular state abundance across longitudinal timepoints was visualized, alongside timepoint pseudotime distributions.

### Bulk RNA-sequencing data gathering and labeling by single-nucleus derived metaprogram

Bulk RNA-sequencing data were obtained from three independent glioma cohorts: The Cancer Genome Atlas (TCGA), the Chinese Glioma Genome Atlas (CGGA) and the Glioma Longitudinal Analysis Consortium (GLASS). For all analyses, we restricted samples to IDH-mutant tumors (including both 1p/19q-codeleted and non-codeleted cases), based on available molecular annotations within each dataset.

For GLASS samples, transcript-level count matrices were downloaded and processed locally. Equivalent raw or normalized count matrices were obtained for TCGA and CGGA. Corresponding clinical and survival annotations were downloaded from each consortium’s associated clinical data releases.

Where applicable, normalization was performed using the EDASeq framework to account for potential within-lane and between-lane technical biases ^153^. Prior to normalization, we evaluated potential GC-content and gene-length bias using diagnostic plots (functions: *biasPlot*, *biasBoxplot*). After confirming no severe systematic technical biases, full offset-based normalization was applied, and normalized counts were extracted.

Defined single-nucleus cellular states were then projected onto bulk RNA-sequencing samples. Briefly, per-sample enrichment testing was assessed using a one-sided Mann-Whitney Wilcoxon gene set test implemented through yaGST (v2017.08.25; https://github.com/miccec/yaGST). Each bulk sample was then assigned a metaprogram according to: (i) the maximum log(NES) value for that sample, (ii) if the associated p-value retained traditional significance (p≤0.05); samples not meeting significance were left as ‘Unassigned’. Metaprogram expression heatmaps were generated using ComplexHeatmap ^154^.

### Kaplan-Meier survival analysis

Overall survival (OS) was defined as the time (in months) from diagnosis to death, or last follow-up. Vital status was encoded as the event (1=deceased, 0=censored), with samples lacking survival time excluded from the analysis. Kaplan-Meier survival curves were estimated using the ‘survival’ package ^155^. Survival distributions were compared across metaprogram-defined cellular states using the log-rank test.

To assess the independent prognostic contribution of cellular state assignment, a multivariable Cox proportional hazards model was fitted including WHO grade, age at diagnosis, and cellular state as covariates. Tie handling was performed using the Efron method. The proportional hazards assumption was evaluated using scaled Schoenfeld residuals. MGMT promoter methylation status was evaluated in a sensitivity analysis, restricted to patients with available data (n=327).

### Clinicopathological stratum enrichment establishment across the bulk RNA-sequencing cohorts

To determine whether specific clinicopathological features were enriched within individual metaprogram-defined groups in the bulk RNA-sequencing datasets, we implemented a custom proportion-based comparison framework.

In short, for each categorical clinicopathological feature, we compared the distribution within each assigned metaprogram grouping against the aggregation of all remaining samples using a two-proportion z-test. Multi-class variables were analyzed by testing each category independently against all other samples. Continuous variables were analyzed through application of a two sided non-parametric Wilcoxon rank-sum test.

### Master Regulator Analysis

Master regulator analysis was performed using DoRothEA, an established and comprehensive gene regulatory network (GRN) resource which catalogs human transcription factors (TFs) and their experimentally or computationally inferred transcriptional targets ^156^. The network was obtained using the *CollecTRI* function, which compiles high-confidence TF-target interactions from multiple databases. Log-normalized count matrices from tumor cell subsets of each sample were used as input and analyzed with the VIPER (Virtual Inference of Protein-activity by Enriched Regulon) algorithm, returning an estimation matrix of normalized enrichment scores (NES) for each TF ^29^. VIPER infers TF activity by evaluating the coordinated expression of a regulator’s target genes rather than the TF itself, yielding a NES for each TF across samples. The resulting activity matrices provided quantitative estimates of TF activity underlying each cellular state. Outputs for each sample were then compiled, and activity estimations were averaged and standard deviation was calculated. Differential activity estimates for TFs were established through non-parametric two-sided Wilcoxon rank-sum tests between activity within and outside the state of interest, with FDR-adjustment. If significantly differential regulators were uniquely positively enriched for a metaprogram, they were highlighted and presented across a ranked-order plot format.

### snATAC-seq Data Processing

Processing and quality control filtration of the snATAC-sequencing (snATAC-seq) dataset was conducted using Signac, a comprehensive package for ATAC-seq data analysis ^157^. Summarily, we generated a unified peak set between all profiled samples to ensure cohort translatability through the package *GenomicRanges* ^158^. Fragment files were then processed on a per-sample basis, and nuclei were initially filtered to remove low-quality barcodes failing basic sequencing and alignment filters (column: ‘passed_filters’).

Quality control metrics were computed for each nucleus, including total ATAC fragment counts, transcription start site (TSS) enrichment scores, fraction of reads in peaks (FRIP), nucleosome banding patterns, and the proportion of reads overlapping genomic blacklist regions. Nuclei were retained using sample-specific and experiment-specific thresholds selected to remove low-complexity or low-signal profiles while preserving biological heterogeneity. Sample specific QC thresholds are detailed in **Supplementary File S1**.

Sequencing depth was summarized using log-transformed ATAC fragment counts, while chromatin signal quality was assessed FRIP and TSS enrichment. The relationship between FRIP and TSS enrichment was evaluated to confirm concordance between global signal-to-noise and promoter-proximal accessibility enrichment. Following filtration on quality control measures, a median of 672 nuclei per sample were retained for downstream analyses.

Samples were normalized, top features (peaks) were selected, and singular value decomposition (SVD) dimension reduction was completed. Nuclei were then projected on a 2-dimensional space (UMAP) using the computed latent semantic indexing (LSI) components, excluding the first component to avoid capturing sequencing depth (technical variation). Cell clusters were identified using LSI components with similar processing, as described in the section ‘**snRNA-seq Data Processing**’.

Extrapolation of cell types captured in snRNA-seq data analyses was conducted through the identification and utilization of reference-based (snRNA-seq) transfer anchors. Anchors were identified using canonical correlation analysis (CCA) on log2-normalized low-dimensional embeddings, following the Seurat label transfer framework ^159^. Each query cell (snATAC-seq) is then assigned a label according to a weighted vote classifier, with each anchor providing a vote weighted by its similarity in behavior to the query cell, as described ^160^. This process was first completed across all cells (malignant and non-tumoral), and then repeated across malignant cells for the informative transfer of labeled cell metaprograms. Low confidence (prediction_score<0.5) cell labels were excluded from downstream analysis.

### snATAC-seq Pseudo-bulking Process

Following identification of malignant cells across the snATAC-seq dataset, downstream pseudo-bulking processes were conducted using the ATAC assay count matrix, representing fragment counts per cell per peak.

Tumor cells were pseudo-bulked per sample by summing accessibility counts and normalizing by cell count. Differential accessibility between GCH and GCL tumors was assessed using edgeR’s glmQLF framework, with peaks at FDR<0.05 considered differentially accessible ^161^. Full CPM normalization rationale and edgeR process detailing may be found in **Supplementary Methods**.

### Motif enrichment analysis of differentially accessible regions

To identify transcription factor (TF) binding motifs enriched within differentially accessible chromatin regions, identified through the pseudobulking process, we implemented Signac ^157^. Human TF position frequency matrices (PFMs) were obtained from the JASPAR 2020 database, restricted to high-confidence, non-redundant motifs ^162^.

Relevant genomic annotations were derived from Ensembl v86 (DOI: <u>10.18129/B9.bioc.EnsDb.Hsapiens.v86</u>) and converted to UCSC-style chromosome naming to ensure compatibility. All analyses were conducted using the hg38 human genome assembly ^163^. Motif occurrences were identified through scanning of accessible chromatin regions using the reference genome. Motifs were then added to the Seurat object, enabling peak-level motif enrichment testing without bias from cell-level sparsity. The identified differentially accessible regions were stratified by directionality of change between GCL and GCH tumors. Motif enrichment was assessed using *FindMotifs*, with p-values adjusted using the Benjamini-Hochberg method. Motifs significantly enriched (top n=20) and congruent with our understanding of GCH and GCL contexts were visualized through motif logo plots.

### Visualization of snATAC-seq derived peaks across Master regulators/putative target genes

Peaks were quantified and normalized using the TF-IDF normalization, followed by selection of highly informative features using peaks with more than 2,750 fragments. Latent semantic indexing (LSI) was performed using singular value decomposition (SVD) with 50 components, and the resulting LSI embeddings were used for visualization and downstream analyses. UMAP was computed using LSI dimensions 2-9. Harmony-based batch correction was evaluated but resulted in reduced separation between GCH and GCL tumor populations, likely reflecting confounding between sample identity and molecular subtype. Unharmonized LSI embeddings were therefore retained for all downstream analyses to preserve biologically relevant chromatin accessibility differences.

Differential accessibility testing was performed separately for each cellular state, comparing cells assigned to a given state against all other tumor cells. Peaks were considered differentially enriched if they satisfied the following criteria: (i) adjusted p-value≤0.05, (ii) higher detection frequency in the state of interest (e.g., pct.1>pct.2), and (iii) a minimum log2 fold-change threshold (state-specific; **Supplementary File S3**). For more stringent analyses focused on putative regulatory interactions, higher fold-change cutoffs were applied. Each peak was assigned to its closest gene using genomic distance to transcription start sites (function: *ClosestFeature*).

Chromatin accessibility at MR and target gene loci was visualized using multi-track coverage plots integrating snATAC-seq accessibility, peak calls, co-accessibility linkages, and matched RNA expression, generated using Signac and Cicero (full visualization parameters available in **Supplementary Methods**) ^157,164^.

### Chromatin co-accessibility analysis (Cicero)

To infer long-range regulatory interactions between distal regulatory elements and gene promoters, chromatin co-accessibility was computed using Cicero (v1.3.9) through generation of co-accessible chromatin accessibility networks (CCANs) ^164^.

### Integrative chromatin analysis: differential histone mark profiling

Chromatin immunoprecipitation sequencing (ChIP-seq) data for H3K27ac and H3K4me3 were analyzed independently using the DiffBind R package (v3.18.0), as mentioned ^146^. Briefly, peak sets were previously generated using MACS2 (v2.2.9.1) and incorporated into DiffBind objects for each mark ^165^. Differential binding analysis comparing GCL versus GCH tumors was performed through DESeq2 implemented within DiffBind ^166^. For each peak, we obtained normalized read concentrations, fold change, alongside nominal and FDR-adjusted p-values. Differential peaks for each mark were annotated to genes using ChIPseeker (v3.22) with the “TxDb.Hsapiens.UCSC.hg38.knownGene” transcript database and gene symbol annotation from org.Hs.eg.db ^167^.

For each histone mark, differential binding statistics were merged with annotated peak coordinates using genomic interval identifiers. To obtain a gene-level summary suitable for integrative analysis, peaks were collapsed on a per-gene basis, selecting the most statistically significant peak associated with each gene. For genes with multiple associated peaks, the peak with the lowest FDR-adjusted p-value was retained as the representative. This strategy captures the strongest evidence of differential chromatin regulation per gene, while avoiding bias from multiple peaks per locus. Genes without peaks for a given mark were assigned neutral values [log2(fold-change)=0; FDR=1].

Mark-specific statistics were then assigned as belonging to: (i) a master regulator (MR), (ii) a regulon member or (iii) unrelated. Statistical significance was defined as FDR≤0.05 for each mark independently. Genes exhibiting significant increases in both H3K27ac (enhancer activation) and H3K4me3 (promoter activation) in GCL relative to GCH tumors (FDR<0.05 for each mark independently) were considered coordinately activated, enabling assessment of chromatin-level regulation associated with MR-driven transcriptional programs.

### Integrative chromatin analysis: genome browser visualization

Genes (MRs and regulons) identified as significant in the differential histone mark-based approach were selected for locus level visualization utilizing both snATAC-seq and ChIP-seq datasets. For each marker, differential H3K27ac and H3K4me3 peaks associated with that gene were extracted, genomic coordinates for the gene body were retrieved using biomaRt (v2.64.0), and a ±25kb window surrounding the gene was defined to provide regulatory context ^168^.

Genome browser tracks were generated using the Gviz package (v1.51.0), with the following tracks constructed: IdeogramTrack, GenomeAxisTrack, BiomartGeneRegionTrack and AnnotationTracks (H3K4me3, H3K27ac, CpG Island indication) ^169^.

snATAC-seq data was incorporated following the aforementioned Signac and Cicero frameworks. Chromatin accessibility coverage plots were generated using Signac’s *CoveragePlot* function ^157^. Coverage was (i) stratified by tumor condition (GCH and GCL), (ii) rendered across the identical genomic window used for ChIP-seq tracks and (iii) extended ±25kb to ensure identical coordinates.

### Immune subsetting and data integration

Upon distinction of the immune component from the rest of the tumor microenvironment, multiple analytical steps were conducted in efforts to functionally stratify these cell types. First, samples were designated to ‘discovery’ (n=8; 1,315 cells) and ‘validation’ (n=6; 172 cells) cohorts based on the number of total non-tumor cells recovered per tumor, with samples contributing ≥100 cells assigned to the discovery cohort, and the remainder to the validation (**Supplementary File S1**). Across the discovery cohort, immune cells from multiple tumor specimens were merged and data preparation and clustering steps were taken, as mentioned (see **Methods** - *snRNA-seq Data Processing*). To mitigate the potential for inter-sample batch effects, integrative non-negative matrix factorization (iNMF) was performed following application of the LIGER (rliger v2.2.1) framework ^17^. Briefly, the RNA assay was split by sample identity, normalized and scaled. iNMF was then performed with k=5 latent factors, followed by quantile normalization to align shared factors across datasets. Clustering and UMAP visualization were then performed using the integrated low-dimensional embedding (*inmfNorm* reduction).

### Compilation of immune reference gene signature sets and correlation with immune subclusters

To infer the identity of immune clusters, curated immune gene signatures were assembled from published brain-specific human immune atlases ^86–88^. Correlative analyses were conducted in which scaled expression values from the LIGER-corrected assays were extracted while excluding mitochondrial genes to minimize technical bias. Expression values were averaged per iNMF-identified cluster to generate a cluster-by-gene matrix, producing a pseudobulk cluster-level expression profile suitable for gene set enrichment comparisons.

To quantify similarity between pseudo-bulked immune clusters and published immune cell types, we applied a rank-based gene set enrichment approach using the Mann-Whitney Wilcoxon gene set test implemented through the yaGST package (v2017.08.25; https://github.com/miccec/yaGST). Briefly, for each cluster: (i) genes were ranked by decreasing mean scaled expression, (ii) for each immune reference signature set, a one-sided Mann-Whitney Wilcoxon test was performed with a minimum gene set size of n=3, and finally (iii) across each cluster-signature set pair, log-transformed enrichment scores and p-values were recorded. Clusters were then “assigned” a putative immune identity if the highest logNES value corresponded to a signature set with p≤0.01.

A heatmap of log-normalized enrichment scores was generated using ComplexHeatmap ^154^. Immune gene programs with at least one positive enrichment score across clusters were retained for visualization. Reference signatures were grouped into broader immune categories (e.g., b-lymphocytes, macrophages, microglial, monocytes, natural killer (NK) and t-lymphocytes), with allowance for hierarchical clustering across both gene programs and immune clusters.

### Construction and characterization of immune ‘super-clusters’

Following consolidation of clusters into six immune super-clusters, we conducted functional analyses through combination of MSigDB and yaGST (v2017.08.25; https://github.com/miccec/yaGST), as described ^170^. Briefly, scaled expression values were averaged across all cells belonging to each cluster to generate pseudo-bulk expression profiles. Genes were then ranked by decreasing mean scaled expression to produce a cluster-level ranked gene list. Functional annotation of immune super-clusters was then performed using gene sets from MSigDB (Hallmark gene sets: H; Curated gene sets: C2; Gene Ontology biological processes: C5), with exclusion for sets unrelated to immune, brain or tumor biology. Rank-based enrichment testing was performed through yaGST framework application (one-sided MWW; function *mwwGST*), with exclusion for gene sets with less than three overlapping genes. To improve interpretability and reduce redundancy, enriched pathways were aggregated based on functional similarity and overlapping gene membership. Redundant terms reflecting the same biological processes were consolidated into higher-level functional themes. Pathways were considered significant at p≤0.01, unless otherwise specified, with visualization of biologically congruent and significant pathways.

### Cytoscape incorporation and network construction

To enable discovery of congruent functional networks implicated in our identified immune super-clusters, we used the pseudobulked expression profiles generated above, in combination with gProfiler2 and Cytoscape (v3.10.3), as described ^147,171^. Briefly, the top 50 genes per super-cluster, based on average expression value, were retained as representative transcriptional programs. Functional enrichment analysis was performed using the gProfiler2 R package across *Homo sapiens*, with corrections for FDR. Networks were constructed in Cytoscape’s EnrichmentMap application, with consolidation of alike-pathways through AutoAnnotate ^149^.

### Utilization of excluded Immune cell populations

To increase the total number of cells attributable to each immune super-cluster, we implemented a targeted labeling approach across immune components which were previously excluded from the initial discovery of cellular states, namely, the validation cohort (see **Supplemental Figure S5a**). Briefly, previously defined immune state transcriptional programs (n=50 genes) were used as reference signatures, and identical yaGST scoring and labeling procedures were applied. Cells were assigned to a super-cluster corresponding to the highest significant normalized enrichment score (NES; p≤0.01). Cell abundance was calculated as the proportion of each cell type, relative to total retrieved immune cells per sample.

### Interpolation of high-CNV immune cells in assessment of clinicopathological dispersion

To further increase the number of cells attributed to each immune super-cluster and more scrupulously discern their relation to clinicopathological features of interest, immune-annotated cells originally excluded for high CNV burden (n=789) were re-evaluated through CNV re-inference and QC metric comparison; those confirmed as transcriptionally equivalent to included immune cells were assigned to the established super-clusters and incorporated into the final immune cohort. Analytical details for the full cell assessment and related visualizations are found in **Supplementary Methods**.

### Cell-cell Communication Analysis

We developed a two-stage ligand-receptor co-expression analysis pipeline inspired by the communication scoring framework of Burdziak et al ^172^. Specifically, we adapted the described quantile bin-profile representation of cellular expression distributions; meanwhile, permutation-based compatibility screenings, graph-theoretic module discovery and fuzzy clustering components represent novel methodological extensions.

Initially, candidate sender-receiver cell population pairs were identified through a permutation-normalized compatibility prescreening across the acquisitioned glioblastoma-derived database of human agonist-receptor based interactions ^173^. Following identification of putative ‘sender’ and ‘receiver’ cell populations, selected pairs were subjected to a directional module detection analysis, which identified co-expressed L-R communication programs specific to the defined candidate populations. Prior to communication analysis, raw UMI count matrices were subjected to Markov Affinity-based Graph Imputation of Cells (MAGIC) imputation to recover gene expression signals characteristic of sparse single-nucleus data ^174^.

### Cell-cell Communication: Stage 1 - Pairwise Compatibility Screening

The initial stage of our communication analysis serves to evaluate ligand-receptor profile compatibility between all possible sender-receiver cell population pairs across G-CIMP strata, producing a ranked interaction matrix which identifies putative communication axes optimal for downstream analyses. First, to account for differences in cell population sizes, across each combination of sample, cell-type and G-CIMP status, log-normalized expression data were represented as quantile bin-profile vectors. Each gene’s expression values across cells within a sample were divided into quantile bins (N=10), and the mean expression within each bin was computed, yielding a genes x10 matrix capturing the distributional shape of expression across the selected cell population. Samples with fewer than two cells of a given population, or fewer than two samples with valid bin profiles per population, were excluded from scoring. Then, for each sender-receiver cell type pair within G-CIMP strata, ligand-receptor compatibility was scored through Pearson correlation calculations using bin-profiles. The observed correlation (ρ) was computed only for ligand-receptor gene pairs where both molecules exceeded an expression threshold of 5% mean expression, consistent with pre-established thresholds, to exclude lowly expressed and likely non-functional interactions ^175^. Per-sample observed scores were averaged across all contributing samples to produce the group-level observed compatibility score.

To establish a null distribution and compute interpretable z-scores, we employed a custom “gene-pair shuffle” null. In each of 200 permutation iterations, the receptor gene indices were randomly assigned among all receptor genes present in the matrix, preserving the bin-profile structure while breaking the biologically meaningful ligand-receptor pairing. This approach produced a null distribution which spans all gene-gene correlations observable in the data, including MAGIC-induced globally elevated correlations, so that resulting z-scores reflect genuine ligand-receptor coordination above values expected from arbitrary gene pairings. Within-strata z-scores were computed as:

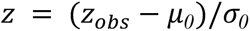

Where the null-mean (*μ_0_*) and standard deviation (*σ_0_*) were estimated across the 200 gene-pair permutations. Resulting z-scores were then stored as a cell-cell compatibility matrix for each G-CIMP group. Sender-receiver pairs with z>0.5 were selected for downstream analysis, a threshold selected to prioritize specificity while retaining sufficient pairings for comprehensive coverage of the tumor microenvironment communication landscape.

### Cell-Cell Communication: Stage 2 - Ligand-Receptor Co-Expression Module Discovery

For each candidate sender-receiver pair, co-expressed ligand-receptor gene modules were identified through a custom pipeline comprising: (2.1) pseudo-bulk bin-profile construction, (2.2) Pearson correlation graph construction with Jaccard similarity pruning, (2.3) PCA-based dimensionality selection using broken-stick and Kneedle elbow criteria, (2.4) spectral eigengap-informed fuzzy c-means clustering, and (2.5) module characterization by gene ontology enrichments. Full algorithmic details can be found in **Supplementary Methods**.

### Cell-Cell Communication: Stage 3 - Automated Pipeline Execution

To ensure comprehensive and systematic coverage of all biologically relevant sender-receiver combinations identified in Stage 1 prescreening, Stage 2’s module discovery was executed through an automated pipeline. This automation eliminates the potential for selective reporting bias, where every interaction exceeding the prescreening z-score threshold (z≥0.5) in either G-CIMP stratum was subjected to the described module protocol.

Following completion of all module discovery runs, per-pair gene ontology (GO) enrichment results were combined into a single aggregated dataset. Semantic similarity-based simplification (GOSemSim v3.22) was applied to results prior to aggregation, collapsing redundant GO terms ^176^. Finally, a cross-pair specificity table was constructed to reveal unique biological specializations of each interaction rather than universal background processes.

### Actionable target identification

To systematically assess the therapeutic tractability of key findings, candidate molecules comprising inferred master regulators, pseudotemporally dynamic genes, ligand-receptor pair components, and enriched pathway effectors were cross referenced with three curated targetable molecule databases: the Drug-Gene Interaction Database (DGIdb), the ChEMBL bioactivity database, and ClinicalTrials.gov [accessed 04/2026]. Candidates were additionally evaluated against a manually curated list of brain-penetrant or CNS-investigated therapeutic agents, defined as compounds with documented blood-brain barrier penetrance, existing CNS clinical trial registration, or prior preclinical evidence in glioma or brain tumor models ^177,178^. Molecules satisfying at least one database interaction criterion and possessing documented or predicted CNS relevance were flagged as prioritized candidates and are discussed in the context of their supporting multi-omic evidence.

### Statistical Analysis

All data processing and analyses were performed in R (v4.5.1). Cell assignment using transcriptional metaprograms was completed either through a combination of one-sided Mann-Whitney Wilcoxon rank-sum test and Seurat’s function Add*ModuleScore*, or through yaGST’s function *mwwGST*. Comparison of cell abundance across two groups was conducted using a two-sided Mann-Whitney Wilcoxon test; across three or more groups using a two-sided Kruskal-Wallis test. Statistical significance for metaprogram enrichment across bulk RNA-sequencing datasets required the maximum log(NES) score and a statistical significance of p≤0.05 (one-sided Mann-Whitney Wilcoxon test), with no applied testing correction. Statistical significance in relation to survival probability using overall survival for stratified metaprogram groupings were established through use of a log-rank test (p≤0.05). Categorical clinicopathological feature enrichments were established using a two-proportion z-test. Continuous feature enrichments were established using a two-sided non-parametric Mann-Whitney Wilcoxon rank-sum test. Correlation analyses were performed using Spearman’s correlation.

### Data Availability

Single-nucleus RNA-sequencing read count data for the acceptable quality malignant and nontumoral cells belonging to samples analyzed throughout this publication have been submitted to Synapse (http://synapse.org; accession #:). Sequestered bulk RNA-sequencing data are available for download from: TCGA (https://portal.gdc.cancer.gov), the CGGA (http://cgga.org.cn/index.jsp) and the GLASS consortium (https://glass-consortium.org/). The multi-omic CGGA dataset used for validation of our single-nucleus findings can be found through dataset ID “CCell_4083” using the aforementioned link. Source data are provided with the paper. All other data supporting the findings of our study are available upon reasonable request from the corresponding author.

### Code Availability

Source codes necessary for the reproduction of all results are available at GitHub [link].

## Acknowledgments

The authors are grateful to the HFH patients who consented to the usage of samples for research purposes. We thank Heather Mengel, Nicolette Bicknell and Lisa Scarpace for their research and clinical administrative support. We thank Ana deCarvalho, Laura Hasselbach and Andrea Transou who obtained patients’ consents, collected and handled the tissue specimens, and maintained the tumor bank at the Hermelin Brain Tumor Center (HBTC). H.N. and A.V.C. are supported by NIH 1R01CA270365-01 and Hermelin Brain Tumor Center and Henry Ford Hospital internal funds.

## Contributions

H.N., A.I. and A.V.C. conceived the study and supervised the analyses and manuscript construction. G.A.H. developed and performed all bioinformatic analyses, requisitioned all data, developed all figures, tables and schematics, and constructed the manuscript under the supervision of H.N., A.I. and A.V.C. Result interpretation and bioinformatic approaches were refined with the help of L.G. and A.I. Multiomic single-nucleus data preparation was completed and detailed by N.S.M. The manuscript was edited and revised by G.A.H., H.N., and A.V.C., with input from all authors.

## Notes

### Competing Interest Statement

The authors have declared no competing interest.

