## Supplementary Information (Methods, Figures, Files) for "Single-nucleus multiomics unveils malignant cellular states, regulatory architectures and microenvironmental reorganization across the G-CIMP epigenomic transition in IDH-mutant glioma": Herrgott et al., Suppl Info.pdf

#### Supplementary Methods

##### *Nuclei isolation protocol details*

*Buffer Preparation*

A 1× Nuclei Buffer was prepared from 20× stock (10x Genomics PN-2000153) containing 1 mM DTT and 1× RNase inhibitor. The lysis buffer contained 10 mM Tris-HCl (pH 7.4), 10 mM NaCl, 3 mM $MgCl_2$, 0.1% Tween-20, 0.1% Nonidet P40 substitute (IGEPAL CA-630), 0.01% Digitonin, 1% BSA, 1 mM DTT, and 1× RNase inhibitor. The wash buffer contained 10 mM Tris-HCl (pH 7.4), 10 mM NaCl, 3 mM $MgCl_2$, 1% BSA, 0.1% Tween-20, 1 mM DTT, and 1× RNase inhibitor.

*Tissue Dissociation and Lysis*

Frozen tissue was mechanically dissociated on ice using a chilled Dounce homogenizer in PBS + 0.04% BSA until a uniform suspension was achieved. The homogenate was filtered through a 40 µm Flowmi cell strainer (Bel-Art H13680-0040) and centrifuged at 300 × g for 5 min at 4 °C. The supernatant was removed, and the pellet was gently resuspended in 100 µL of chilled lysis buffer. Samples were incubated on ice for 3–5 min to lyse cellular membranes, with incubation time optimized to yield intact, free nuclei without cytoplasmic debris.

*Washing and Resuspension*

Following lysis, 1 mL of chilled wash buffer was added, and the suspension was gently mixed by pipetting 5×. Nuclei were pelleted through centrifugation at 500 × g for 5 min at 4 °C, and the supernatant was carefully removed without disturbing the pellet. This wash step was repeated twice for a total of three washes.

After the final wash, nuclei were resuspended in 1× Diluted Nuclei Buffer (10x Genomics) and maintained on ice. The Tris/$Mg^{2+}$ composition of this buffer is optimized for compatibility with downstream transposition and barcoding steps of the Chromium Next GEM Single Cell Multiome ATAC + GEX workflow.

##### *Nuclei Preparation, Quality Control, and Loading*

Isolated nuclei suspensions were filtered through a 40 µm Flowmi cell strainer when debris or clumping was observed. Nuclei concentration and integrity were evaluated using a hemocytometer on an EVOS XL Core Imaging System following 0.4% Trypan Blue staining. Samples exhibiting predominantly intact nuclei with minimal debris were considered suitable for downstream processing. Nuclei were counted and diluted in 1× Nuclei Buffer to the target concentration in accordance with 10x Genomics guidelines (typically 3,000–10,000 nuclei per reaction), and maintained on ice to preserve nuclear integrity. A calculated number of nuclei were then loaded onto a Chromium Next GEM Chip J, together with reagents for simultaneous ATAC transposition and mRNA capture, and all subsequent steps - including GEM generation and library construction - were performed according to the manufacturer`s protocol (Chromium Next GEM Single Cell Multiome ATAC + Gene Expression User Guide, CG000338).

##### *Longitudinal Pseudotime Validation Across Matched Primary and Recurrent Samples*

To assess whether pseudotime dynamics were preserved across longitudinal disease progression, we performed a patient-specific trajectory validation analysis using four patients for whom matched primary and recurrent specimens were available. Six of the ten samples were present in the pooled trajectory model cohort; the remaining four samples had been previously excluded due to insufficient malignant cell recovery (n<250 cells), but were retained for this validation. For samples present in the pooled model, scaled pseudotime values were extracted directly from the Slingshot trajectory analysis as described above. For the excluded samples, pseudotime was estimated via reference-based projection onto the pooled Harmony embedding. Briefly, each sample's scaled expression values were projected onto the pooled PCA space by matrix multiplication against the pooled PCA rotation matrix, yielding per-cell coordinates in the 25-dimensional PCA space. Harmony correction was then applied to a combined matrix of reference (pooled) and query (excluded sample) PCA embeddings, using sample identity as the grouping variable. Corrected coordinates for query cells were extracted, pseudotime values were assigned to each cell via inverse-distance-weighted k-nearest-neighbor interpolation (k=15) in the 15-dimensional Harmony embedding space, using the reference cells' known pseudotime values as the interpolation target. Projection quality was assessed by computing the concordance between each projected cell's assigned metaprogram label and the majority metaprogram label among its 15 nearest reference neighbors; samples with concordance <0.5 were flagged for exclusion, though all four samples exceeded this threshold.

***snATAC-seq Pseudo-bulking Process***

To improve statistical robustness and mitigate the potential for interference of characteristic sparsity in chromatin accessibility data, a pseudo-bulking strategy was employed. Rather than aggregating across conditions, accessibility counts were summed across cells within each sample, preserving biological replication and avoiding pseudoreplication which would arise from treating individual cells as independent observations. Then, due to the variable attribution of each sample towards total cell count, counts were normalized by the number of contributing cells per sample. Specifically, summed peak counts were divided by the total number of cells in each sample, yielding a CPM-like normalization which reflects accessibility per cell rather than total sequencing depth alone. To assess regions for differential accessibility trends between G-CIMP High and Low tumors, we applied the edgeR framework [1]. Briefly, low-abundance peaks were filtered using *filterByExpr*, requiring sufficient counts across biological replicates. Differential accessibility testing was conducted using a generalized linear model quasi-likelihood F-test (glmQLF) due to its robustness in experiments with heterogeneous variability. Peaks with FDR-adjusted p-value$\leq$0.05 were considered differentially accessible. For visualization purposes, normalized values were log2-transformed after addition of a negligible pseudocount.

*Visualization of snATAC-seq derived peaks across Master regulators/putative target genes*

For selected master regulators and putative target genes, genomic regions of interest (ROIs) were defined based on differentially accessible peaks proximal to or overlapping the gene locus. For each ROI, multi-track coverage plots were generated using Signac, integrating the following layers of regulatory information: (i) chromatin accessibility coverage tracks, stratified by cellular state; (ii) tile plots highlighting accessibility specifically within the state of interest; (iii) peak annotation tracks; (iv) gene annotation tracks displaying gene models and TSS; (v) Cicero co-accessibility links [2,3]. Additionally, RNA expression plots were generated from the complementary RNA assay, grouped by cellular state, to assess concordance between chromatin accessibility and transcriptional output.

*Interpolation of high-CNV immune cells in assessment of clinicopathological dispersion*

Cells originally annotated as 'Immune system cells' or 'Microglial cells' through our scType application, and excluded for high CNV burden, were extracted (n=789). To ensure that excluded cells exhibited true CNV signal rather than technical noise, we repeated our CNV-inference pipeline, where: (i) a non-denoised inferCNV analysis was rerun, (ii) CNV signal intensity and correlation to reference cells were quantified, (iii) density distributions of CNV signal were compared between previously included and excluded cells.

To determine whether elevated CNV burden across immune populations was associated with poor cell quality, standard QC metrics were evaluated, including: total UMI counts, number of features, percent mitochondrial and percent ribosomal gene expression. Features were compared between previously included and excluded cells to ensure equivalent distribution of quality.

To align high-CNV cells with previously defined immune super-clusters, identical labeling processes were reapplied using the transcriptional programs (n=50 genes), as detailed. Cells with no associated significant relations were labeled as 'Unknown'.

Finally, high-CNV labeled immune cells were combined with: (i) previously included (discovery) immune cells, and (ii) independently labeled excluded cells to generate a comprehensive picture of the total immune component for our cohort. To assess cell distributions across tumor contexts, proportional abundance per tumor sample was calculated. Associations between cell abundance (per sample) and the following clinicopathological features was assessed: G-CIMP status, tumor grade, time of collection (initial or recurrent). Wilcoxon rank-sum tests were applied to compare subtype abundance distributions between groups, with significant differences highlighted.

***Cell-Cell Communication Stage 2***

*2.1 Expression matrix construction*

For each candidate sender-receiver pair identified in Stage 1, a detailed communication module discovery pipeline was applied to identify the co-expressed ligand-receptor gene programs which delineate communication between populations.

Firstly, depending on population size, those cell types possessing >15 cells were compiled into a pseudo-bulk bin-profile representation, constructed through division of cells into 15 quantile bins per gene per population, yielding 30 pseudo-observations. Ligand-receptor genes were then pre-filtered to those expressed above the 50th percentile mean expression in the designated population, retaining pairs where at least one direction met this threshold. Genes with near-zero variance ($<10^{-6}$) were removed.

*2.2 Gene-Gene Correlation Graph Construction*

A Pearson correlation matrix was then computed across all retained ligand and receptor genes using the constructed matrix. An initial adjacency matrix was created through thresholding correlations ($\rho<0.01$), and a weighted undirected graph was constructed. To remove spurious co-expression edges which arise from global expression trends, Jaccard similarity pruning was applied. Edges with a similarity below the 10th percentile of all non-zero Jaccard values were removed, and isolated nodes were excluded.

*2.3 Dimensionality Reduction: Broken-Stick and Geometric Elbow Criterion*

Principal component analysis (PCA) was applied to the pruned gene-gene correlation matrix to identify the major axes of co-variation across modules. The number of informative PCs was

determined by the joint application of (i) a broken stick model and (ii) the geometric elbow ("Kneedle" method).
Briefly, the broken stick model compares the observed variance explained by each component against the expectation under a null of randomly distributed variance. Components whose variance exceeds the expected value are retained. The geometric elbow, or "Kneedle" method, identifies the point of maximum curvature on the scree curve. Most frequently, this is the component where the curve transitions from steep signal-containing decline to a flat noise-denominated tail [4]. In calculations, both axes of the scree are normalized before computing the perpendicular distance from the chord connecting the initial and culminatory points, and the component with the maximum distance is designated as the "elbow".
We enforced a minimum of 2 and maximum of 30 components; additionally, the within-cluster variance scan for module count estimation was constrained so that the number of candidate cluster centers never exceeded the number of selected components, preventing degenerate fuzzy clusterings.

*2.4 Module Count Estimation & Assignment*

The optimal number of communication modules was estimated in two steps: first, the spectral eigengap of the normalized graph Laplacian was computed on the pruned correlation graph (2.2), identifying the number of clusters (k) where the gap between consecutive eigenvalues was maximized. Second, a within-cluster variance scan was conducted across a window of $\pm 4$ candidate k-values around the spectral estimate, with the second derivative of the within-cluster variance curve used to identify the elbow of the WCV plot as the final module count.
Within-cluster variance was computed using Fuzzy c-means (m=2.0) and Euclidean distance in the PC space [5].
Module assignment was conducted through Fuzzy c-means clustering application across the PC-score matrix using the selected number of modules. This yielded a continuous membership probability matrix for the selected ligand and receptor molecules. Genes were assigned to modules with membership probability exceeding 0.2, a threshold selected to permit biologically meaningful bridge membership while excluding genes with only marginal association to a given module. Modules with less than three member genes were discarded.

*2.5 Module Characterization and Enrichment*

Each module was annotated with ligand and receptor role assignments based on membership in the L-R annotation database. Mean expression values per gene in the sender and receiver cell populations were computed to rank the most active genes within each module. Per-module interaction figures, generated in Biorender.com, were informed upon bipartite sender-receiver graphs, complete with edge width and opacity measures (data not shown).
Ontological biological processes (BP), molecular functions (MF) and cellular components (CC) enrichment analysis were performed for each module using clusterProfiler (v4) [6]. Term enrichment significance was assessed using the hypergeometric test with Benjamini-Hochberg correction.

**Supplementary File Descriptions**

**Supplementary File S1 -** Single-nucleus sample clinicopathological and quality control information

**Supplementary File S2 -** Malignant cell relevant signature sets

**Supplementary File S3 -** Multi-OMIC validation effort information

**Supplementary File S4 -** Immune cell signatures and cell-cell communication information

**Supplementary Figure Legends**

**Supplementary Figure S1. a)** Histograms of filtered cells across several quality features including: (i) total counts, (ii) number of detected genes, (iii) mitochondrial and (iv) ribosomal expression proportion. Vertical lines represent 10th and 90th percentiles**. b)** Boxplots illustrating distribution of copy number aberration (i) correlation and (ii) signal, per cell. **c)** Uniform manifold approximation and projection (UMAP) for dimension reduction of 4,112 non-tumoral cells based on gene expression. **d)** Scatterplots depicting the estimated enrichment of published cell type expression programs across non-tumor clusters. Y-axis represents the log-scaled significance (p-value), x-axis represents the log-scaled normalized enrichment scores (NES). **e)** Boxplot displaying cell type abundance across samples, split by (i) combinatory G-CIMP and Time of Collection (ii) G-CIMP status, and (iii) WHO Grade. **Note**: *: p<0.05; **: p<0.01, ***: p<0.001; ****: p<0.0001.

**Supplementary Figure S2. a)** Uniform manifold approximation and projection (UMAP) for dimension reduction based on gene expression of the 10,335 malignant cells. **b)** Similarity heatmaps denoting assignment of published cellular state and internal cellular state assignments. **c)** Correlation heatmaps denoting the relation of cellular states across (i) module scoring and (ii) variant gene expression. **d)** Abundance bar plots displaying cell cycle phase membership across each malignant cellular state. **e)** EnrichmentMap networks for the identified malignant cell states, representing organized significant ontological terms. **f)** Histogram displaying the distribution of cells stratified by program of interest across cell complexity. Vertical lines represent 10th and 90th percentiles. **g)** Positively scored and differentially enriched Transcription factor (TF) activity heatmap, with columnar hierarchical clustering.

**Supplementary Figure S3. a)** Rank order plot displaying neuronal signature set expression correlation scores across cellular states. **b)** Dot plot displaying expression-values across cell groups, split by G-CIMP. **c)** Boxplot displaying the distribution of normalized enrichment scores possessed by GABA-OPC tumor cells across Oligodendrocyte, oligodendrocyte precursor (OPC), and GABA-OPC hybrid signature sets. **d)** Uniform manifold approximation and projections (UMAPs) for dimension reduction based on gene expression of 7,128 cells from malignant and nontumoral populations, colored by G-CIMP. (Bottom) Cells are colored by expression levels of two OPC hallmark markers. **e-f)** Hierarchical clusterings of functional terms enriched for upregulated markers derived through the comparisons of: **(e)** GABA-OPC malignant and non-tumor oligodendrocyte, and **(f)** Neu-NPC malignant and non-tumor neuronal

(InN, ExN). **g)** UMAPs for dimension reduction based on gene expression of 9,626 malignant cells. Cells are colored by: (i) cellular states, and (ii) scaled pseudotime. Notes: GCL: G-CIMP-low; GCH: G-CIMP-high.

**Supplementary Figure S4. (a-d)** A series of plots including: (i) scaled-pseudotime boxplots, segregated by collection and colored by G-CIMP; (ii) Uniform manifold approximation and projections (UMAP) for dimensionality reduction per patient, colored by pseudotime, and highlighted by sample; (iii) stacked barplots displaying cell abundance of cellular states, per collection. Series are repeated and labeled for each longitudinal patient.

**Supplementary Figure S5. a-c)** Expression heatmaps displaying metaprograms (MPs) across bulk samples sequestered from **(a)** CGGA, **(b)** TCGA and **(c)** GLASS data sources. Samples are annotated with relevant clinicopathological or molecular features; genes are grouped by MP. **d)** Barplot displaying abundance of cellular states across CGGA single-cell data. Columns are annotated with relevant clinicopathological information, and clustered by assigned state. **e)** (left) Promoter-region CpG sites hypomethylation barplots focused in the state of interest. (right) Scatterplots displaying the relation of protein abundance and gene expression. Pearson correlation statistics are included. Related density plots for dispersion of points across each axis are included. Note: *: $p<0.05$; **: $p<0.01$, ***: $p<0.001$; ****: $p<0.0001$. **f)** Scatterplot depicting relation of master regulator (MR) and regulon expression, associated with the particular cellular state of interest. **g)** Quality control plot series for snATAC-seq data including: (i) snATAC-seq counts, (ii) fraction of reads in peaks (FRiP), and (iii) relation between transcription start site (TSS) enrichment and FRiP, grouped by G-CIMP. Pearson correlation statistics are displayed. Horizontal/vertical lines annotate 10th and 90th percentiles. **h-i)** Genome-browser tracks visualizing the presence of ChIP-seq and snATAC-seq results across selected loci.

**Supplementary Figure S6. a)** Immune-based analytical workflow. **b)** Uniform Manifold Approximation and Projection (UMAP) for dimension reduction across 1,315 immune cells taken, grouped by cluster. **c)** Scatter plots depicting the enrichment of selected pathways. The y-axis represents log-scaled significance (p-value), x-axis represents the normalized enrichment scores. **d)** Density plot displaying the estimated copy number signal for both included and excluded cells. Vertical lines represent median signal values. **e)** Histograms displaying the distribution of immune cells across quality features including: (i) total counts, (ii) number of detected genes, (iii) mitochondrial and (iv) ribosomal expression proportion. Vertical lines represent 10th and 90th percentiles. **f)** Heatmap depicting module expression scores of immune super-clusters across high-CNV cells. g) Boxplot displaying super-cluster abundance across samples, split by (i) G-CIMP, and (ii) WHO Grade. Note: *: $p<0.05$; **: $p<0.01$, ***: $p<0.001$; ****: $p<0.0001$.

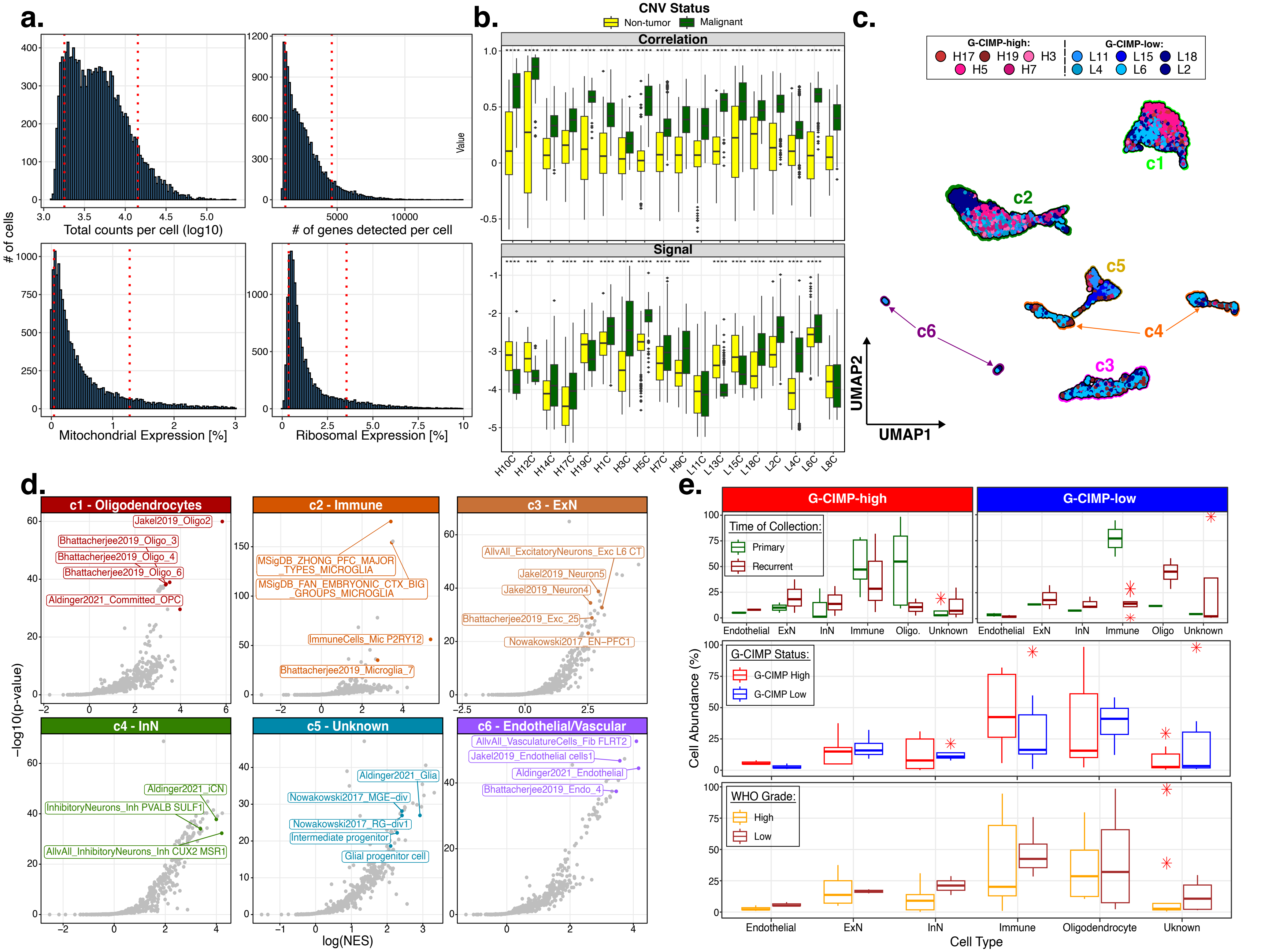
a.
### of cells
Total counts per cell (log10)
### of genes detected per cell
Mitochondrial Expression [%]
Ribosomal Expression [%]
b.
CNV Status
Non-tumor
Malignant
Correlation
Signal
Value
H10C H12C H14C H17C H19C H1C H3C H5C H7C H9C L11C L13C L15C L18C L2C L4C L6C L8C
c.
G-CIMP-high:
H17 H19 H3
H5 H7
G-CIMP-low:
L11 L15 L18
L4 L6 L2
c1
c2
c3
c4
c5
c6
UMAP1
UMAP2
d.
c1 - Oligodendrocytes
Jakel2019_Oligo2
Bhattacherjee2019_Oligo_3
Bhattacherjee2019_Oligo_4
Bhattacherjee2019_Oligo_6
Aldinger2021_Committed_OPC
c2 - Immune
MSigDB_ZHONG_PFC_MAJOR_TYPES_MICROGLIA
MSigDB_FAN_EMBRYONIC_CTX_BIG_GROUPS_MICROGLIA
ImmuneCells_Mic P2RY12
Bhattacherjee2019_Microglia_7
c3 - ExN
AllvAll_ExcitatoryNeurons_Exc L6 CT
Jakel2019_Neuron5
Jakel2019_Neuron4
Bhattacherjee2019_Exc_25
Nowakowski2017_EN−PFC1
c4 - InN
Aldinger2021_iCN
InhibitoryNeurons_Inh PVALB SULF1
AllvAll_InhibitoryNeurons_Inh CUX2 MSR1
c5 - Unknown
Aldinger2021_Glia
Nowakowski2017_MGE−div
Nowakowski2017_RG−div1
Intermediate progenitor
Glial progenitor cell
c6 - Endothelial/Vascular
AllvAll_VasculatureCells_Fib FLRT2
Jakel2019_Endothelial cells1
Aldinger2021_Endothelial
Bhattacherjee2019_Endo_4
−log10(p-value)
log(NES)
e.
G-CIMP-high
G-CIMP-low
Time of Collection:
Primary
Recurrent
Endothelial ExN InN Immune Oligo. Unknown
Endothelial ExN InN Immune Oligo Unknown
G-CIMP Status:
G-CIMP High
G-CIMP Low
WHO Grade:
High
Low
Cell Abundance (%)
Endothelial ExN InN Immune Oligodendrocyte Uknown
Cell Type

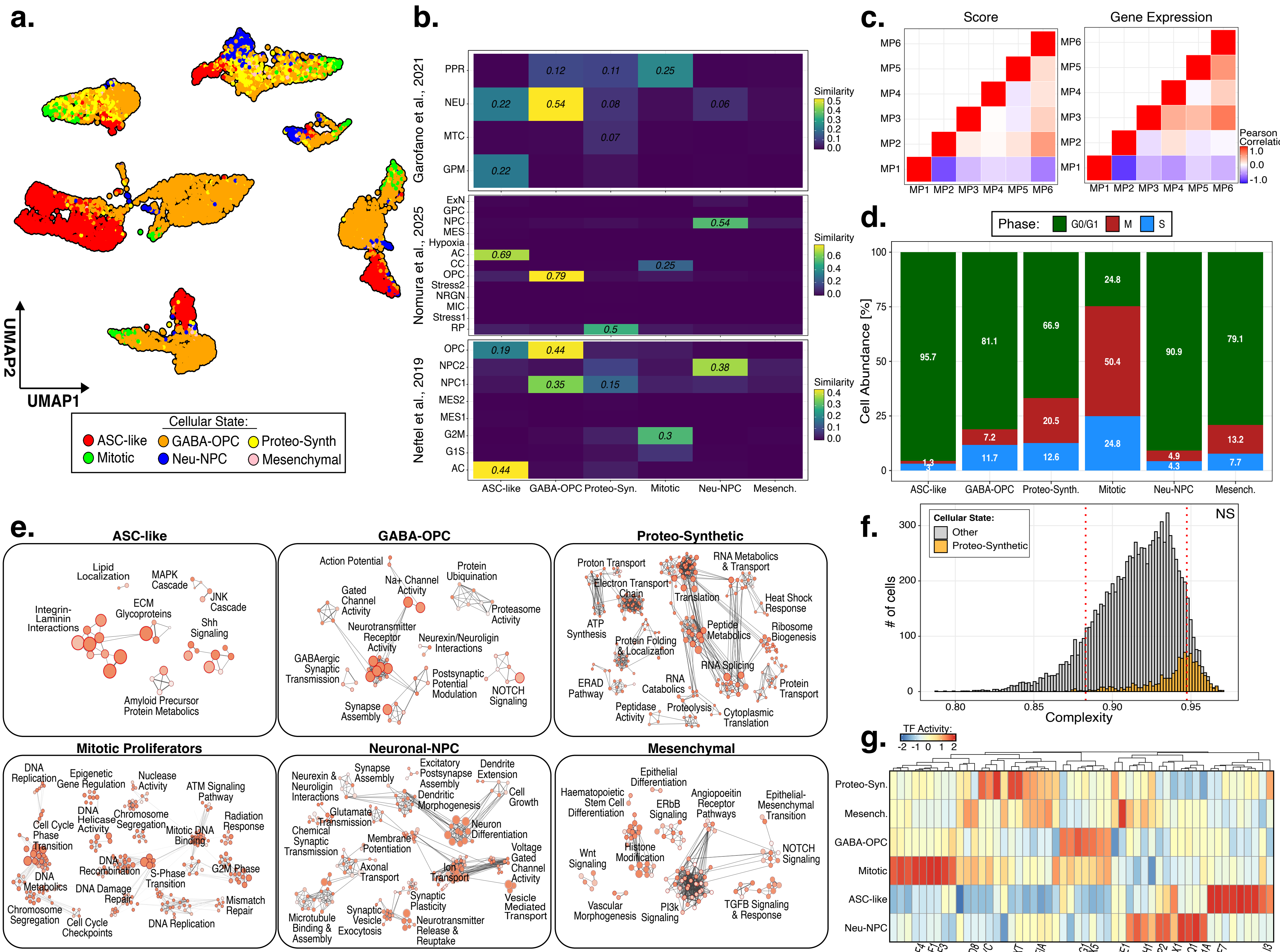
a.
UMAP2
UMAP1
Cellular State:
ASC-like
GABA-OPC
Proteo-Synth
Mitotic
Neu-NPC
Mesenchymal
b.
Garofano et al., 2021
PPR
NEU
MTC
GPM
0.12
0.11
0.25
0.22
0.54
0.08
0.06
0.07
0.22
Similarity
Nomura et al., 2025
ExN
GPC
NPC
MES
Hypoxia
AC
CC
OPC
Stress2
NRGN
MIC
Stress1
RP
0.54
0.69
0.25
0.79
0.5
Neftel et al., 2019
OPC
NPC2
NPC1
MES2
MES1
G2M
G1S
AC
0.19
0.44
0.38
0.35
0.15
0.3
0.44
ASC-like
GABA-OPC
Proteo-Syn.
Mitotic
Neu-NPC
Mesench.
c.
Score
Gene Expression
MP1
MP2
MP3
MP4
MP5
MP6
Pearson Correlation
1.0
0.0
-1.0
d.
Phase:
G0/G1
M
S
Cell Abundance [%]
95.7
1.3
3
81.1
7.2
11.7
66.9
20.5
12.6
24.8
50.4
24.8
90.9
4.9
4.3
79.1
13.2
7.7
ASC-like
GABA-OPC
Proteo-Synth.
Mitotic
Neu-NPC
Mesench.
e.
ASC-like
Lipid Localization
MAPK Cascade
JNK Cascade
ECM Glycoproteins
Integrin-Laminin Interactions
Shh Signaling
Amyloid Precursor Protein Metabolics
GABA-OPC
Action Potential
Protein Ubiquination
Na+ Channel Activity
Gated Channel Activity
Proteasome Activity
Neurotransmitter Receptor Activity
Neurexin/Neuroligin Interactions
GABAergic Synaptic Transmission
Postsynaptic Potential Modulation
NOTCH Signaling
Synapse Assembly
Proteo-Synthetic
Proton Transport
RNA Metabolics & Transport
Electron Transport Chain
Translation
Heat Shock Response
ATP Synthesis
Peptide Metabolics
Ribosome Biogenesis
Protein Folding & Localization
RNA Splicing
ERAD Pathway
RNA Catabolics
Protein Transport
Peptidase Activity
Proteolysis
Cytoplasmic Translation
Mitotic Proliferators
DNA Replication
Epigenetic Gene Regulation
Nuclease Activity
ATM Signaling Pathway
Cell Cycle Phase Transition
DNA Helicase Activity
Chromosome Segregation
Mitotic DNA Binding
Radiation Response
DNA Recombination
S-Phase Transition
G2M Phase
DNA Metabolics
DNA Damage Repair
Mismatch Repair
Chromosome Segregation
Cell Cycle Checkpoints
DNA Replication
Neuronal-NPC
Neurexin & Neuroligin Interactions
Synapse Assembly
Excitatory Postsynapse Assembly
Dendrite Extension
Dendritic Morphogenesis
Cell Growth
Glutamate Transmission
Chemical Synaptic Transmission
Membrane Potentiation
Neuron Differentiation
Voltage Gated Channel Activity
Axonal Transport
Ion Transport
Synaptic Plasticity
Synaptic Vesicle Exocytosis
Microtubule Binding & Assembly
Neurotransmitter Release & Reuptake
Vesicle Mediated Transport
Mesenchymal
Epithelial Differentiation
Haematopoietic Stem Cell Differentiation
ERbB Signaling
Angiopoeitin Receptor Pathways
Epithelial-Mesenchymal Transition
Histone Modification
NOTCH Signaling
Wnt Signaling
Vascular Morphogenesis
PI3k Signaling
TGFB Signaling & Response
f.
Cellular State:
Other
Proteo-Synthetic
NS
### of cells
Complexity
g.
TF Activity:
Proteo-Syn.
Mesench.
GABA-OPC
Mitotic
ASC-like
Neu-NPC
E2F4
E2F1
E2F3
HOXD8
MYC
TBXT
NFIA
OLIG1
TBX5
FOXE1
NHLH1
FOXP2
DLX1
FOXQ1
BCL11A
KLF7
GLI3

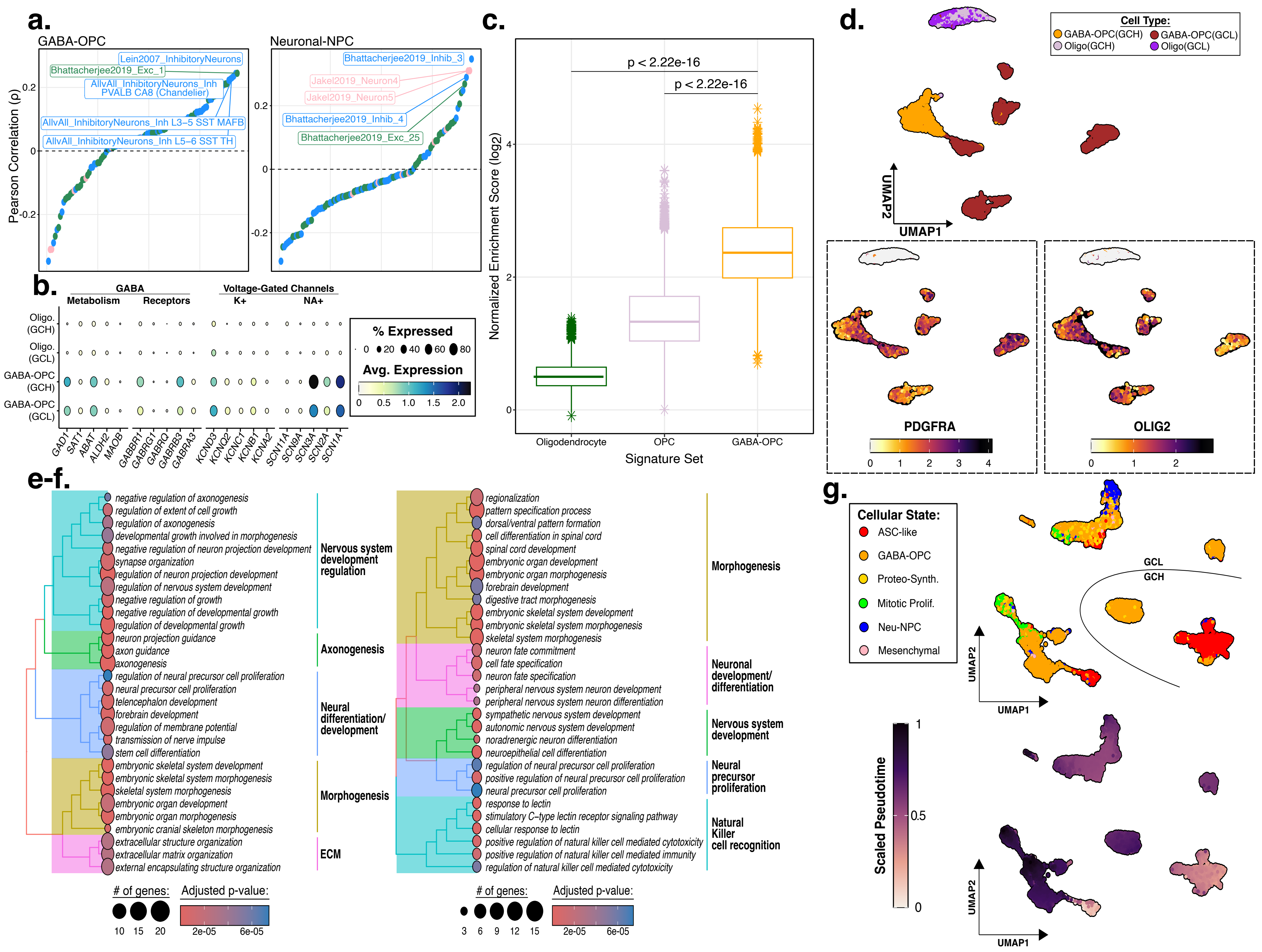

a.
GABA-OPC
Neuronal-NPC
Pearson Correlation (ρ)
Lein2007_InhibitoryNeurons
Bhattacherjee2019_Exc_1
AllvAll_InhibitoryNeurons_Inh PVALB CA8 (Chandelier)
AllvAll_InhibitoryNeurons_Inh L3−5 SST MAFB
AllvAll_InhibitoryNeurons_Inh L5−6 SST TH
Bhattacherjee2019_Inhib_3
Jakel2019_Neuron4
Jakel2019_Neuron5
Bhattacherjee2019_Inhib_4
Bhattacherjee2019_Exc_25
b.
GABA
Metabolism
Receptors
Voltage-Gated Channels
K+
NA+
Oligo. (GCH)
Oligo. (GCL)
GABA-OPC (GCH)
GABA-OPC (GCL)
% Expressed
Avg. Expression
GAD1 SAT1 ABAT ALDH2 MAOB GABBR1 GABRG1 GABRQ GABRB3 GABRA3 KCND3 KCNQ2 KCNC1 KCNB1 KCNA2 SCN11A SCN9A SCN3A SCN2A SCN1A
c.
p < 2.22e-16
p < 2.22e-16
Normalized Enrichment Score (log2)
Oligodendrocyte
OPC
GABA-OPC
Signature Set
d.
Cell Type:
GABA-OPC(GCH)
GABA-OPC(GCL)
Oligo(GCH)
Oligo(GCL)
UMAP1
UMAP2
PDGFRA
OLIG2
e-f.
negative regulation of axonogenesis
regulation of extent of cell growth
regulation of axonogenesis
developmental growth involved in morphogenesis
negative regulation of neuron projection development
synapse organization
regulation of neuron projection development
regulation of nervous system development
negative regulation of growth
negative regulation of developmental growth
regulation of developmental growth
neuron projection guidance
axon guidance
axonogenesis
regulation of neural precursor cell proliferation
neural precursor cell proliferation
telencephalon development
forebrain development
regulation of membrane potential
transmission of nerve impulse
stem cell differentiation
embryonic skeletal system development
embryonic skeletal system morphogenesis
skeletal system morphogenesis
embryonic organ development
embryonic organ morphogenesis
embryonic cranial skeleton morphogenesis
extracellular structure organization
extracellular matrix organization
external encapsulating structure organization
Nervous system development regulation
Axonogenesis
Neural differentiation/ development
Morphogenesis
ECM
regionalization
pattern specification process
dorsal/ventral pattern formation
cell differentiation in spinal cord
spinal cord development
embryonic organ development
embryonic organ morphogenesis
forebrain development
digestive tract morphogenesis
embryonic skeletal system development
embryonic skeletal system morphogenesis
skeletal system morphogenesis
neuron fate commitment
cell fate specification
neuron fate specification
peripheral nervous system neuron development
peripheral nervous system neuron differentiation
sympathetic nervous system development
autonomic nervous system development
noradrenergic neuron differentiation
neuroepithelial cell differentiation
regulation of neural precursor cell proliferation
positive regulation of neural precursor cell proliferation
neural precursor cell proliferation
response to lectin
stimulatory C−type lectin receptor signaling pathway
cellular response to lectin
positive regulation of natural killer cell mediated cytotoxicity
positive regulation of natural killer cell mediated immunity
regulation of natural killer cell mediated cytotoxicity
Morphogenesis
Neuronal development/ differentiation
Nervous system development
Neural precursor proliferation
Natural Killer cell recognition
### of genes:
Adjusted p-value:
g.
Cellular State:
ASC-like
GABA-OPC
Proteo-Synth.
Mitotic Prolif.
Neu-NPC
Mesenchymal
GCL
GCH
UMAP1
UMAP2
Scaled Pseudotime

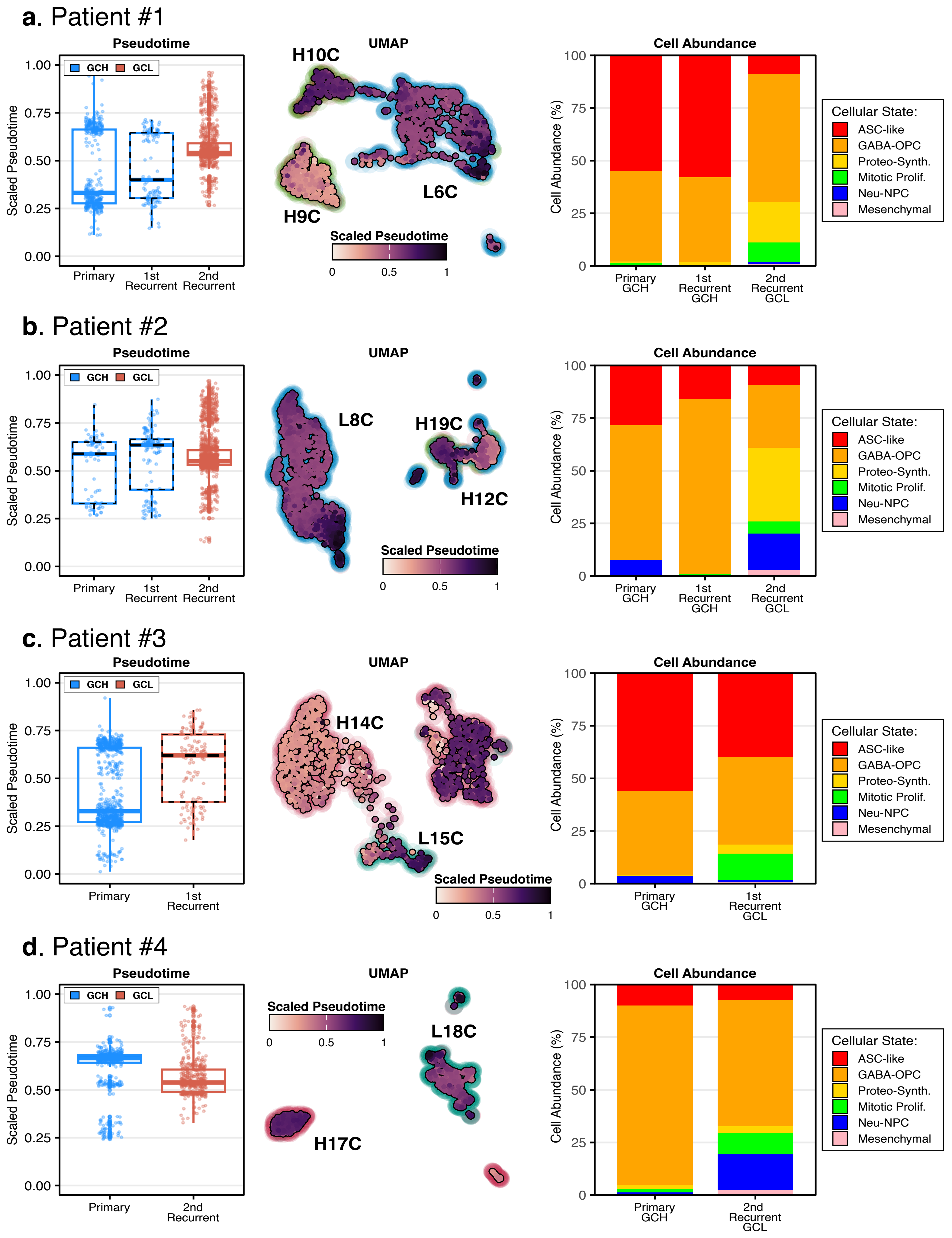
a. Patient #1
Pseudotime
GCH
GCL
Scaled Pseudotime
1.00
0.75
0.50
0.25
0.00
Primary
1st Recurrent
2nd Recurrent
UMAP
H10C
H9C
L6C
Scaled Pseudotime
0
0.5
1
Cell Abundance
Cell Abundance (%)
100
75
50
25
0
Primary GCH
1st Recurrent GCH
2nd Recurrent GCL
Cellular State:
ASC-like
GABA-OPC
Proteo-Synth.
Mitotic Prolif.
Neu-NPC
Mesenchymal
b. Patient #2
Pseudotime
GCH
GCL
Scaled Pseudotime
Primary
1st Recurrent
2nd Recurrent
UMAP
L8C
H19C
H12C
Scaled Pseudotime
Cell Abundance
Cell Abundance (%)
Primary GCH
1st Recurrent GCH
2nd Recurrent GCL
Cellular State:
ASC-like
GABA-OPC
Proteo-Synth.
Mitotic Prolif.
Neu-NPC
Mesenchymal
c. Patient #3
Pseudotime
GCH
GCL
Scaled Pseudotime
Primary
1st Recurrent
UMAP
H14C
L15C
Scaled Pseudotime
Cell Abundance
Cell Abundance (%)
Primary GCH
1st Recurrent GCL
Cellular State:
ASC-like
GABA-OPC
Proteo-Synth.
Mitotic Prolif.
Neu-NPC
Mesenchymal
d. Patient #4
Pseudotime
GCH
GCL
Scaled Pseudotime
Primary
2nd Recurrent
UMAP
Scaled Pseudotime
L18C
H17C
Cell Abundance
Cell Abundance (%)
Primary GCH
2nd Recurrent GCL
Cellular State:
ASC-like
GABA-OPC
Proteo-Synth.
Mitotic Prolif.
Neu-NPC
Mesenchymal

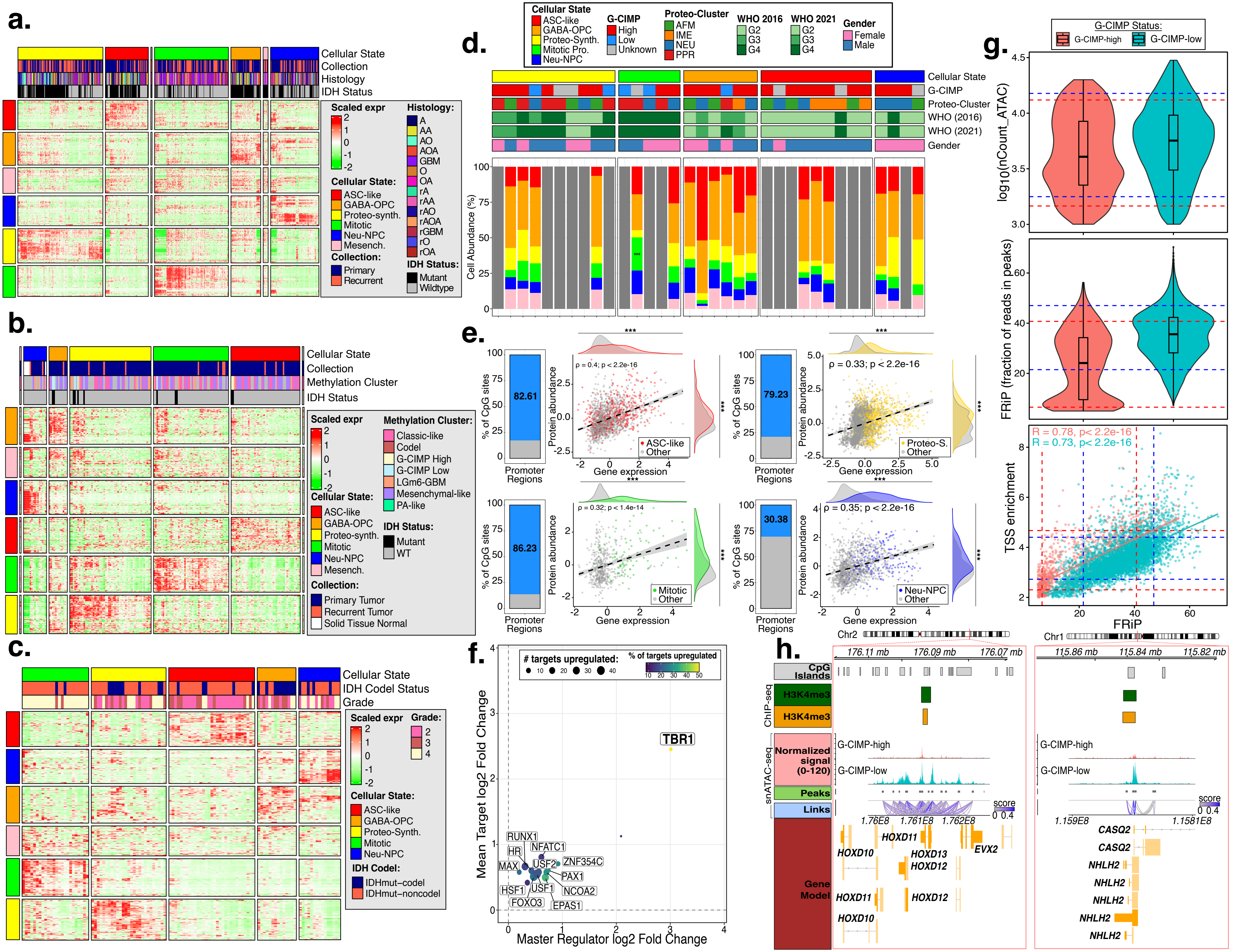

a.
Cellular State
Collection
Histology
IDH Status
Scaled expr
Histology:
A
AA
AO
AOA
GBM
O
OA
rA
rAA
rAO
rAOA
rGBM
rO
rOA
Cellular State:
ASC-like
GABA-OPC
Proteo-synth.
Mitotic
Neu-NPC
Mesench.
Collection:
Primary
Recurrent
IDH Status:
Mutant
Wildtype
b.
Cellular State
Collection
Methylation Cluster
IDH Status
Scaled expr
Methylation Cluster:
Classic-like
Codel
G-CIMP High
G-CIMP Low
LGm6-GBM
Mesenchymal-like
PA-like
Cellular State:
ASC-like
GABA-OPC
Proteo-synth.
Mitotic
Neu-NPC
Mesench.
IDH Status:
Mutant
WT
Collection:
Primary Tumor
Recurrent Tumor
Solid Tissue Normal
c.
Cellular State
IDH Codel Status
Grade
Scaled expr
Grade:
2
3
4
Cellular State:
ASC-like
GABA-OPC
Proteo-Synth.
Mitotic
Neu-NPC
IDH Codel:
IDHmut–codel
IDHmut–noncodel
d.
Cellular State
ASC-like
GABA-OPC
Proteo-Synth.
Mitotic Pro.
Neu-NPC
G-CIMP
High
Low
Unknown
Proteo-Cluster
AFM
IME
NEU
PPR
WHO 2016
G2
G3
G4
WHO 2021
G2
G3
G4
Gender
Female
Male
Cellular State
G-CIMP
Proteo-Cluster
WHO (2016)
WHO (2021)
Gender
Cell Abundance (%)
e.
% of CpG sites
Promoter Regions
82.61
79.23
86.23
30.38
Protein abundance
Gene expression
ρ = 0.4; p < 2.2e-16
ρ = 0.33; p < 2.2e-16
ρ = 0.32; p < 1.4e-14
ρ = 0.35; p < 2.2e-16
ASC-like
Other
Proteo-S.
Other
Mitotic
Other
Neu-NPC
Other
f.
### targets upregulated:
% of targets upregulated
TBR1
RUNX1
HR
NFATC1
MAX
USF2
ZNF354C
PAX1
HSF1
USF1
NCOA2
FOXO3
EPAS1
Mean Target log2 Fold Change
Master Regulator log2 Fold Change
g.
G-CIMP Status:
G-CIMP-high
G-CIMP-low
log10(nCount_ATAC)
FRiP (fraction of reads in peaks)
R = 0.78, p< 2.2e-16
R = 0.73, p< 2.2e-16
TSS enrichment
FRiP
h.
Chr2
Chr1
176.11 mb
176.09 mb
176.07 mb
115.86 mb
115.84 mb
115.82 mb
CpG Islands
ChIP-seq
H3K4me3
H3K4me3
snATAC-seq
Normalized signal (0-120)
G-CIMP-high
G-CIMP-low
Peaks
Links
score
1.76E8
1.761E8
1.762E8
1.159E8
1.1581E8
Gene Model
HOXD11
HOXD10
HOXD13
HOXD12
EVX2
CASQ2
NHLH2

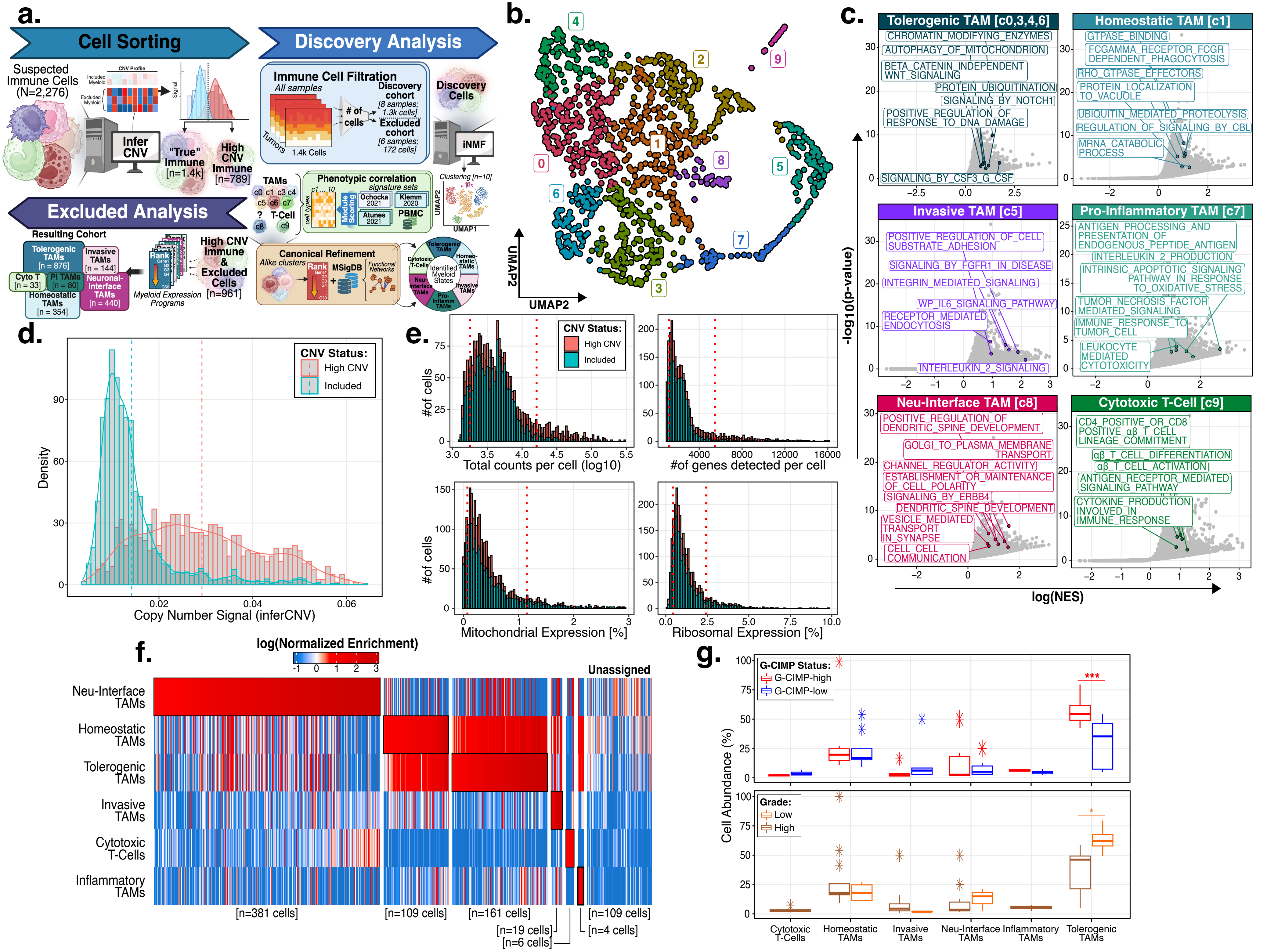

a.
Cell Sorting
Suspected Immune Cells (N=2,276)
CNV Profile
Infer CNV
"True" Immune [n=1.4k]
High CNV Immune [n=789]
Discovery Analysis
Immune Cell Filtration
All samples
Discovery cohort [8 samples; 1.3k cells]
### of cells
Excluded cohort [6 samples; 172 cells]
Tumors
1.4k Cells
Discovery Cells
iNMF
Clustering [n=10]
UMAP1
UMAP2
TAMs
c0 c1 c3 c4 c5 c6 c7
T-Cell
c8 c9
Phenotypic correlation
signature sets
Module Scoring
Ochocka 2021
Klemm 2020
Atunes 2021
PBMC
Canonical Refinement
Alike clusters
Rank
MSigDB
Functional Networks
Identified Myeloid States
Tolerogenic TAMs
Homeostatic TAMs
Invasive TAMs
Pro-Inflamm TAMs
Neu-Interface TAMs
Cytotoxic T-Cells
Excluded Analysis
Resulting Cohort
Tolerogenic TAMs [n = 876]
Invasive TAMs [n = 144]
Cyto T [n = 33]
PI TAMs [n = 80]
Neuronal-Interface TAMs [n = 440]
Homeostatic TAMs [n = 354]
Myeloid Expression Programs
High CNV Immune & Excluded Cells [n=961]
b.
UMAP1
UMAP2
0 1 2 3 4 5 6 7 8 9
c.
Tolerogenic TAM [c0,3,4,6]
CHROMATIN_MODIFYING_ENZYMES
AUTOPHAGY_OF_MITOCHONDRION
BETA_CATENIN_INDEPENDENT WNT_SIGNALING
PROTEIN_UBIQUITINATION
SIGNALING_BY_NOTCH1
POSITIVE_REGULATION_OF RESPONSE_TO_DNA_DAMAGE
SIGNALING_BY_CSF3_G_CSF
Homeostatic TAM [c1]
GTPASE_BINDING
FCGAMMA_RECEPTOR_FCGR DEPENDENT_PHAGOCYTOSIS
RHO_GTPASE_EFFECTORS
PROTEIN_LOCALIZATION TO_VACUOLE
UBIQUITIN_MEDIATED_PROTEOLYSIS
REGULATION_OF_SIGNALING_BY_CBL
MRNA_CATABOLIC PROCESS
Invasive TAM [c5]
POSITIVE_REGULATION_OF_CELL SUBSTRATE_ADHESION
SIGNALING_BY_FGFR1_IN_DISEASE
INTEGRIN_MEDIATED_SIGNALING
WP_IL6_SIGNALING_PATHWAY
RECEPTOR_MEDIATED ENDOCYTOSIS
INTERLEUKIN_2_SIGNALING
Pro-Inflammatory TAM [c7]
ANTIGEN_PROCESSING_AND PRESENTATION_OF ENDOGENOUS_PEPTIDE_ANTIGEN
INTERLEUKIN_2_PRODUCTION
INTRINSIC_APOPTOTIC_SIGNALING PATHWAY_IN_RESPONSE TO_OXIDATIVE_STRESS
TUMOR_NECROSIS_FACTOR MEDIATED_SIGNALING
IMMUNE_RESPONSE_TO TUMOR_CELL
LEUKOCYTE MEDIATED CYTOTOXICITY
Neu-Interface TAM [c8]
POSITIVE_REGULATION_OF DENDRITIC_SPINE_DEVELOPMENT
GOLGI_TO_PLASMA_MEMBRANE TRANSPORT
CHANNEL_REGULATOR_ACTIVITY
ESTABLISHMENT_OR_MAINTENANCE OF_CELL_POLARITY
SIGNALING_BY_ERBB4
DENDRITIC_SPINE_DEVELOPMENT
VESICLE_MEDIATED TRANSPORT IN_SYNAPSE
CELL_CELL COMMUNICATION
Cytotoxic T-Cell [c9]
CD4_POSITIVE_OR_CD8 POSITIVE_αβ_T_CELL LINEAGE_COMMITMENT
αβ_T_CELL_DIFFERENTIATION
αβ_T_CELL_ACTIVATION
ANTIGEN_RECEPTOR_MEDIATED SIGNALING_PATHWAY
CYTOKINE_PRODUCTION INVOLVED_IN IMMUNE_RESPONSE
-log10(p-value)
log(NES)
d.
CNV Status:
High CNV
Included
Density
Copy Number Signal (inferCNV)
e.
CNV Status:
High CNV
Included
#of cells
Total counts per cell (log10)
#of genes detected per cell
Mitochondrial Expression [%]
Ribosomal Expression [%]
f.
log(Normalized Enrichment)
Unassigned
Neu-Interface TAMs
Homeostatic TAMs
Tolerogenic TAMs
Invasive TAMs
Cytotoxic T-Cells
Inflammatory TAMs
[n=381 cells]
[n=109 cells]
[n=161 cells]
[n=19 cells]
[n=6 cells]
[n=109 cells]
[n=4 cells]
g.
G-CIMP Status:
G-CIMP-high
G-CIMP-low
***
Grade:
Low
High
*
Cell Abundance (%)
Cytotoxic T-Cells
Homeostatic TAMs
Invasive TAMs
Neu-Interface TAMs
Inflammatory TAMs
Tolerogenic TAMs
